# VEFill: a model for accurate and generalizable deep mutational scanning score imputation across protein domains

**DOI:** 10.1101/2025.05.14.653991

**Authors:** Polina V Polunina, Wolfgang Maier, Alan F Rubin

## Abstract

**Background:** Deep Mutational Scanning (DMS) assays can systematically assess the effects of amino acid substitutions on protein function. While DMS datasets have been generated for many targets, they often suffer from incomplete variant coverage due to technical constraints, limiting their utility in variant interpretation and downstream analyses.

**Results:** We developed VEFill, a gradient boosting model for imputing missing DMS scores across protein domains. VEFill is trained on the Human Domainome 1 dataset, a large, standardized set of DMS experiments using a uniform stability-based assay, and integrates a broad set of additional biologically informative features including ESM-1v sequence embeddings, evolutionary conservation (EVE scores), amino acid substitution matrices, and physicochemical descriptors. The model achieved robust predictive performance (***R***^***2***^ = 0.64, Pearson r = 0.80). It also demonstrated reliable generalization to unseen proteins in other stability-based datasets, while showing weaker performance on activity-based assays. Per-protein models further confirmed VEFill’s effectiveness under limited-data conditions. A reduced two-feature version using only ESM-1v embeddings and mean DMS scores performed comparably to the full model, suggesting a computationally efficient alternative. However, true zeroshot prediction without positional context remains a challenge, particularly for functionally complex proteins.

**Conclusions:** VEFill offers an interpretable, scalable framework for DMS score imputation, especially effective in stability-focused and sparse-data settings. It enables systematic mutation prioritization and may support the design of efficient experimental libraries for variant effect studies.

## 1 Introduction

Deep mutational scanning (DMS) is a high-throughput experimental technique that systematically assays the functional consequences of thousands of protein variants in parallel, generating detailed variant effect (VE) maps [1, 2]. These maps are essential for elucidating protein structure–function relationships, predicting the impact of missense mutations, and interpreting genetic variation in both research and clinical contexts. As of 2025, MaveDB [3]—a curated repository for multiplexed assays of variant effects (MAVEs)—lists over 2,600 publicly available datasets, including more than 1,100 for human proteins. Although the technology is advancing rapidly, most DMS datasets remain incomplete due to experimental limitations such as low coverage, sequencing depth constraints, and assay dropout, leaving many mutations without experimental scores. This missing data hinders large-scale integrative analyses and downstream applications in variant prioritization and protein engineering.

Several supervised and unsupervised models have been developed to impute missing values in DMS datasets by leveraging evolutionary, biochemical, or structural features. Random forest-based approaches [4, 5] use averaged or position-weighted features including BLOSUM62 scores [6], and physicochemical differences, achieving reasonable accuracy on small numbers of curated proteins. Gradient boosting models like Envision [7] extended this framework with additional conservation and structural metrics. Deep learning methods, such as Deep2Full [8], have incorporated neural networks with sequence and co-evolutionary features, while FUSE [9] introduced a shrinkage estimation pipeline using James-Stein-derived position-level means. Other recent methods like AALasso and FactorizeDMS [10] have explored variant similarity and latent matrix decomposition, respectively. Other approaches focus on imputation based solely on DMS scores without incorporating external features. For example, a recent method imputes missing scores by identifying the median of the most similar amino acid substitutions based on BLOSUM100 similarity within the same protein context, followed by refinement using weighted averaging with observed values [11, 12]. While these models vary in feature sets and complexity, most are either limited in scalability, restricted to a few datasets, or tailored to specific proteins or assay types.

Despite growing interest in DMS score imputation, existing models often have limited generalizability, due to being trained on small or protein-specific datasets, or relying on features that are not uniformly available across proteins, such as structural annotations [4, 7, 8, 10]. Imputation approaches may also be optimized for individual proteins [4, 5] and primarily leverage substitution matrices (e.g., BLOSUM62) or conservation-based scores (e.g., SIFT [13], PROVEAN [14]). While effective in certain contexts, these features largely capture broad evolutionary constraints and may fail to reflect more complex dependencies, such as long-range residue interactions, that influence mutational effects. Recent work on CPT-1 [15]—a cross-protein transfer learning model for clinical variant classification—demonstrated strong performance across genes by combining features such as EVE scores [16], amino acid properties, and embeddings from ESM-1v—a transformer-based language model trained on protein sequences that captures long-range dependencies via its attention mechanism [17]. Although CPT-1 can be adapted for other tasks, it is primarily designed to classify variants as pathogenic or benign. This highlights a remaining need for a scalable and robust framework tailored specifically to the task of DMS score imputation.

To address this gap, we developed VEFill, a scalable and generalizable model for DMS score imputation that integrates diverse biologically informed features. Trained on the Human Domainome 1 dataset [18]—a large, standardized dataset based on a uniform stability assay—VEFill is designed to generalize across unseen proteins and variant types without relying on structural annotations (Fig. 1). In addition to the DMS data, the model incorporates EVE scores as evolutionary constraints, amino acid physicochemical properties, amino acid substitution matrices, and ESM-1v sequence embeddings, the latter of which are designed to capture long-range dependencies within the amino acid sequence and improve predictions in a context-sensitive manner.

**Fig. 1:**
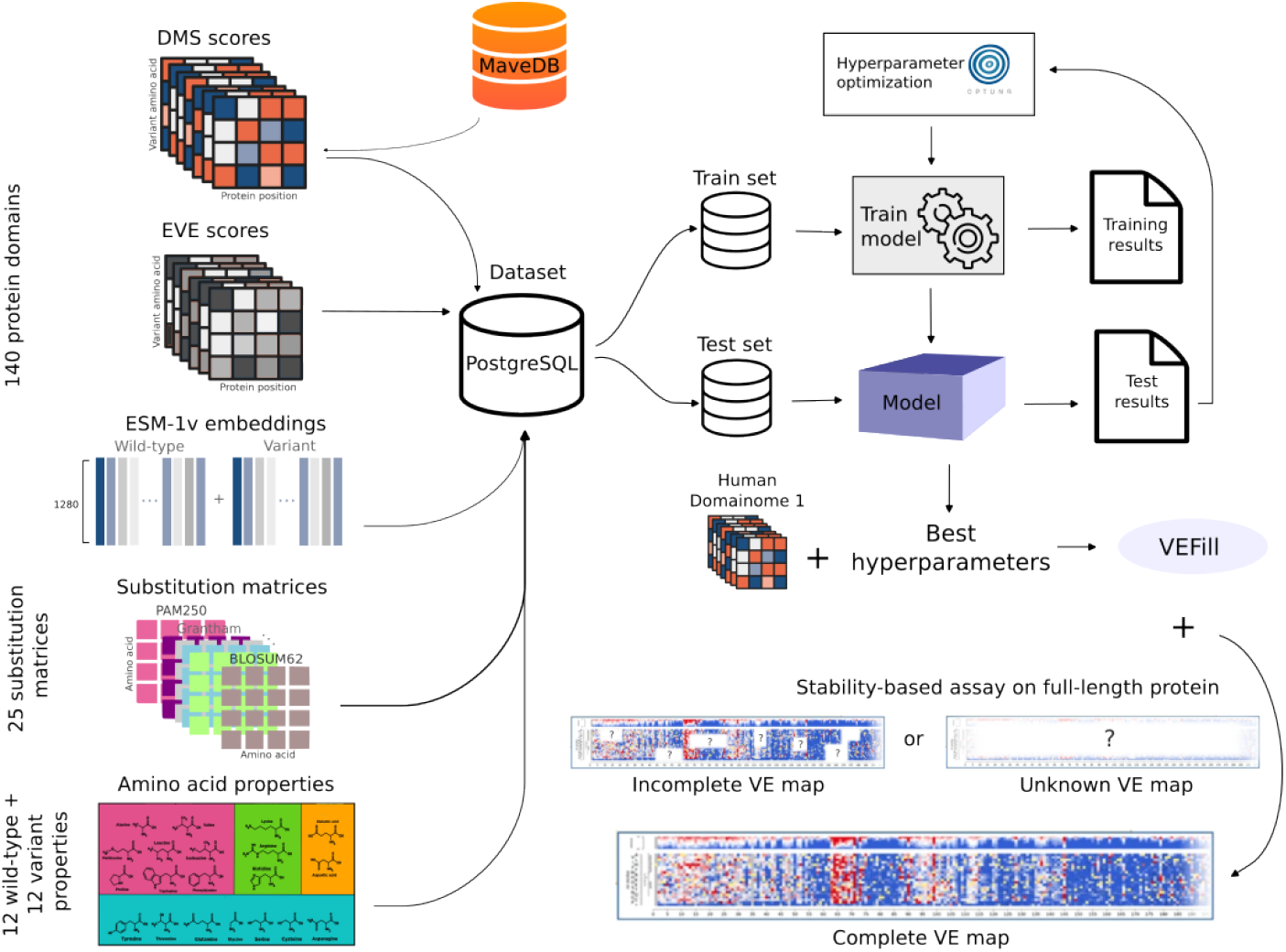
VEFill model design, optimization, training, and application. Features collected include experimental DMS scores from human protein domains (obtained from MaveDB), EVE scores, ESM-1v embeddings, amino acid substitution matrices, and various physicochemical amino acid properties. All datasets required for model training, testing, and inference were stored in a PostgreSQL database with a custom schema. A LightGBM gradient-boosting model was trained and optimized using Bayesian hyperparameter optimization (via Optuna). The trained VEFill model can effectively impute missing DMS scores and predict variant effects, including for stability-based assays on previously unseen full-length proteins in a zero-shot manner.

## 2 Methods

### 2.1 Data collection and storage

For this study, we collected DMS datasets from MaveDB, focusing on the Human Domainome 1 dataset, which originally includes mutagenesis data for 522 protein domains. One of three artificial (non-natural) sequences from the original Human Domainome 1 dataset was excluded due to its absence in MaveDB, resulting in 521 domains available for downstream analysis. This decision ensured consistency in data sourcing and prioritized biologically grounded sequences.

For general model development, we used a filtered subset of 140 domains for which EVE scores were available. This filtering step ensured consistent feature representation across all proteins and avoided introducing data sparsity or inconsistencies in downstream modeling. The full set of 521 domains was used to train reduced-feature models relying only on ESM-1v embeddings and mean DMS scores, simulating settings where fewer feature types are available (see Supplementary Table S1 for inclusion details).

EVE scores were retrieved from the official EVE model website. A total of 25 amino acid substitution matrices were obtained—24 from the `Biopython v1.83` package [19] (via `Bio.Align.substitution_matrices`) and one from a publicly available source (Supplementary Table S2). Physicochemical properties were derived from established biochemical literature and databases. Features such as hydrophobicity, solvent accessibility, charge, chemical group classifications, molecular weight, pKa values, isoelectric point, hydropathy index, and others were obtained from curated biochemical sources, including the AAindex database [20], Kyte-Doolittle hydropathy scale [21], and Lehninger Principles of Biochemistry [22] (see Supplementary Table S3 for the full list of features used for VEFill, including physicochemical properties).

Protein embeddings were generated using the protein language model ESM-1v (650M parameters, UR90S/1 variant; model identifier: `esm1v_t33_650M_UR90S_1.porgt`) [17]. The pre-trained model weights were downloaded from the FAIR ESM repository (https://github.com/facebookresearch/esm). For each protein, we retrieved residue-level embeddings for both the wild-type sequence and corresponding single-amino-acid variants from the model’s final layer (layer 33), utilizing the official `fair-esm` Python package (version 2.0.0). Additionally, we computed difference vectors between wildtype and variant embeddings at the mutated positions to specifically capture localized sequence changes.

To reconcile different indexing schemes—MaveDB reports positions relative to the domain (1-based), whereas EVE uses UniProt [23] full-length coordinates—we identified offset values for each protein to allow for correct mapping during preprocessing.

All collected data, including target amino acid sequences, UniProt offsets, DMS scores, EVE scores, substitution matrices, embeddings, and other features, were stored in a `PostgreSQL v14.17` database for structured, scalable, and unified access throughout the modeling pipeline (see Supplementary Fig. S1 for a schema of the database architecture).

### 2.2 Feature selection

To capture the multifactorial nature of mutational effects, we integrated diverse feature types into our model. These were selected to reflect evolutionary constraints, biochemical properties, and sequence context.

Substitution matrices (e.g., BLOSUM62, PAM250 [24]) provide general evolutionary-informed scores for amino acid replacements. Physicochemical properties (e.g., charge, polarity, volume, hydrophobicity) were included for wild-type, variant, and difference values to reflect structural and functional impacts. A single-nucleotide variant (SNV) indicator feature was included to distinguish more naturally likely substitutions from multi-nucleotide changes.

We incorporated ESM-1v embeddings to capture contextual, learned representations of protein sequences, and EVE scores to reflect evolutionary conservation and summarize the information from multiple sequence alignments. Additionally, we included the mean DMS score per position as a data-driven prior to account for site-specific mutational tolerance.

This feature set was designed to balance biological interpretability with high predictive performance (see Supplementary Table S3 for a complete list of features).

### 2.3 Model architecture

We selected LightGBM [25], a gradient boosting framework optimized for efficiency and scalability, as the core model for VEFill. The task involves structured, high-dimensional tabular data and requires regression-based modeling, making tree-based ensembles particularly suitable. LightGBM offers strong predictive performance on medium-to-large datasets without the need for deep learning–scale data volumes. Its built-in regularization and leaf-wise tree growth help prevent overfitting, while support for GPU acceleration ensures computational efficiency. Compared to fully connected neural networks (a classic multilayer perceptrons) [26] or transformer-based models [27], LightGBM achieved competitive accuracy in our preliminary tests with substantially faster training and easier interpretability, making it a pragmatic and effective choice for this study (Supplementary Table S4).

### 2.4 Data normalization and preprocessing

Raw DMS scores were normalized using Eq. 1, where 0 corresponds to the median score of nonsense mutations (loss-of-function baseline) and 1 corresponds to the wild-type score. The normalized DMS score *s*_*norm*_ was computed as:

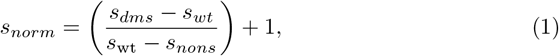

where *s*_*dms*_ is the raw DMS score of the variant, *s*_*wt*_ is the wild-type score determined by the DMS study, and *s*_*nons*_ is the mean score of nonsense mutations determined by the DMS study.

This transformation centers neutral mutations (similar to WT) around 1, while deleterious variants typically fall near 0. Some variants—particularly hyperstable mutations or scoring artifacts—can yield normalized scores slightly above 1 or below 0, reflecting that normalization preserves the relative scale of raw measurements and does not bound scores strictly to the [0, 1] interval.

All features were preprocessed to ensure compatibility with the LightGBM model. Categorical variables—such as amino acid class descriptors (e.g., chemical group, charge)—were one-hot encoded and included for wild-type and variant amino acids. Boolean biochemical descriptors (e.g., hydrogen bonding capability, solvent accessibility) were converted to binary integers (0 or 1). Both boolean (converted to binary) and numerical properties of amino acids (e.g., molecular weight, isoelectric point, pKa, hydropathy index) were included for the wild-type and variant residues, as well as their absolute difference:

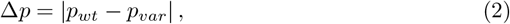

where *p* denotes the respective amino acid property, *p*_*wt*_ and *p*_*var*_ are the values of the property for the wild-type and variant amino acids, respectively.

ESM-1v embeddings for wild-type, variant, and difference vectors were flattened into fixed-length numeric arrays and appended as individual input features. The difference vector Δ**e** was computed as:

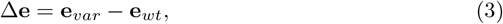

where **e**_*wt*_ and **e**_*var*_ denote the embedding vectors at the mutated position in the wild-type and variant sequences, respectively. This operation captures the local contextual shift in the learned representation space induced by the mutation.

Additionally, we computed the mean normalized DMS score per residue position, grouped by protein and position index. This feature served as a data-driven prior reflecting mutational tolerance at each site, and was used as an additional input feature. The position-wise mean was computed as:

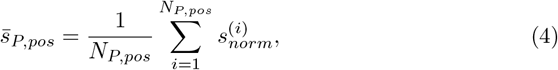

where 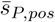 is the mean normalized DMS score for protein *P* at position *pos*, 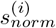 is the normalized score of the *i*^*th*^ variant at that position, and 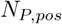 is the total number of variants observed at position *pos* in protein *P*.

All preprocessing steps were executed using a Python-based pipeline that integrated SQL querying from a `PostgreSQL v14.17` database, followed by transformation using `pandas v2.2.3` [28, 29] and `NumPy v2.2.0` [30]. This ensured reproducibility and consistent data formatting across the dataset.

### 2.5 Hyperparameter optimization

To optimize model performance, we performed hyperparameter tuning using the Optuna framework [31], which applies Bayesian optimization through a Tree-structured Parzen Estimator (TPE) sampler [32]. A group-aware 5-fold cross-validation strategy was used, with protein domains (identified by gene IDs from our custom PostgreSQL database) serving as grouping variables to prevent data leakage. The objective was to minimize the root mean squared error (RMSE) across folds.

The LightGBM model was trained using gradient boosting decision trees with up to 1,000 boosting rounds and early stopping triggered after 50 consecutive rounds without improvement. The search space covered key hyperparameters including learning rate, maximum depth, number of leaves, subsampling ratios, and regularization strengths (L1 and L2 penalties). Once the optimal configuration was identified, a final model was trained using the selected parameters and evaluated on a held-out 10% test set, stratified by gene ID. Evaluation metrics included RMSE, mean absolute error (MAE), and the coefficient of determination (*R*^2^).

### 2.6 Model variants and training strategies

To evaluate imputation strategies for DMS scores, we trained several variants of LightGBM-based models using different data partitioning schemes and feature compositions. All models shared the common LightGBM architecture and were trained using optimized hyperparameters.

#### 2.6.1 General model

The general model was trained on 136,854 mutations spanning 140 protein domains from the Human Domainome 1 dataset. A leave-protein-out split was applied, where 90% of the protein domains were used for training and the remaining 10% were held out as a test set. To investigate the contribution of different types of biological information, we trained the general model using a range of feature set configurations combining evolutionary (EVE scores, substitution matrices), biochemical (amino acid properties), sequence-derived contextual (ESM-1v embeddings), and position-aware (mean DMS score per position, SNV indicator) features. These configurations are summarized in Fig. 2 and detailed in Supplementary Fig. S2 and Supplementary Table S5, where model performance metrics (*R*^2^, Pearson r, RMSE, and MAE) are compared across feature combinations. We observed major performance gains upon inclusion of ESM-1v embeddings, with further improvement when mean DMS scores per position were added.

**Fig. 2:**
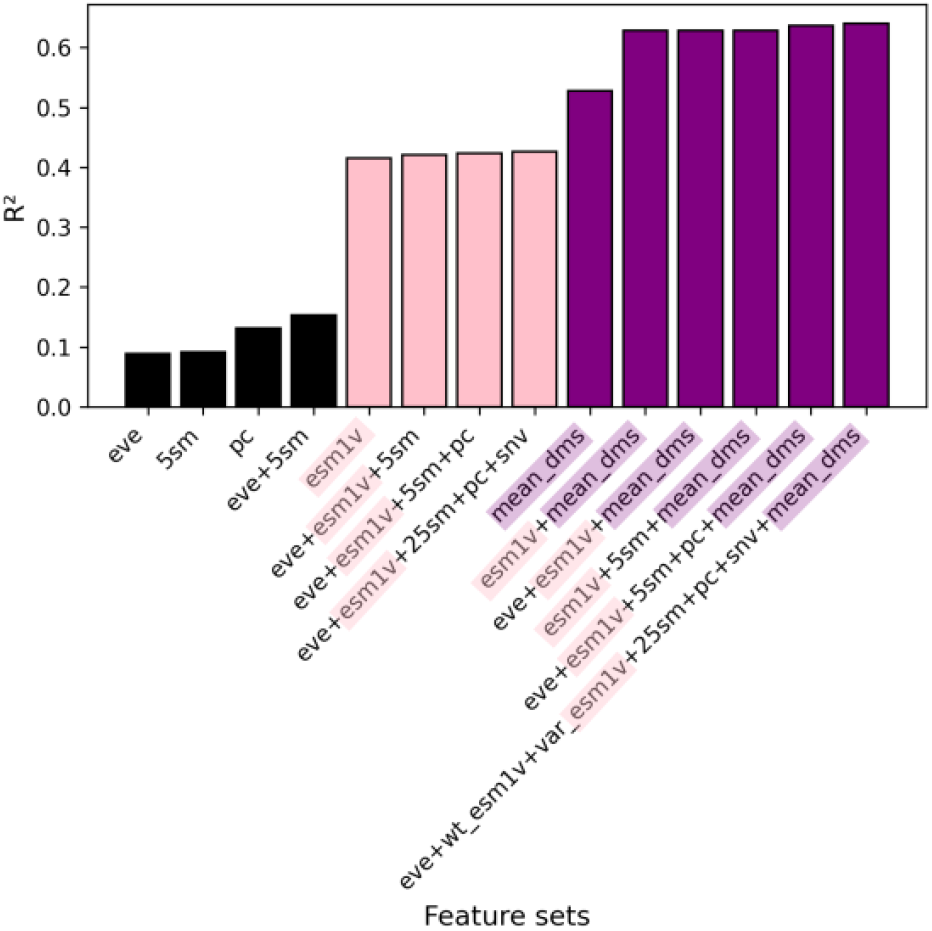
General model performance (*R*^2^) for DMS score imputation across feature sets. Models were trained on the 140 protein domains for which EVE scores were available. Predictive power increased with feature richness, especially when ESM-1v embeddings and positional priors (mean DMS scores) were included. The top model (*R*^2^ = 0.64, r = 0.80) excluded only the ESM-1v difference vector (see Supplementary Table S3 for full definitions of feature sets, and Supplementary Fig. S2 and Table S5 for full results).

#### 2.6.2 General (LOPO) model

For a more systematic evaluation of model generalizability, we implemented a leave-one-protein-out (LOPO) cross-validation strategy. In this setup, 140 models were trained, each time holding out one protein domain as the test set while training on the remaining 139 domains.

#### 2.6.3 Per-protein models

To benchmark localized prediction performance, we trained per-protein models individually for each of the 140 protein domains using several train/test partitioning strategies. In the random split setup, each protein’s mutation dataset was randomly divided into training and test subsets using various ratios, including 90/10, 85/15, 80/20, 20/80, and 15/85. In the leave-position-out (LPosO) configuration, positions within each protein were split into training and test sets based on 80/20 and 20/80 ratios, allowing the model to generalize across sequence regions. The leave-one-position-out (LOPosO) approach involved holding out all variants at a single position as the test set while training on variants from the remaining positions. Lastly, in the leave-one-variant-out (LOVarO) strategy, each variant was sequentially held out for testing, with all other variants in the same protein used for training.

In addition to using the full feature set, per-protein models were also trained with reduced feature configurations informed by feature importance analyses from the general model. These included a reduced set containing only ESM-1v embeddings and the mean DMS score per position, as well as a set containing all features except ESM-1v embeddings.

#### 2.6.4 Minimal feature model

To evaluate the impact of training dataset size and feature sparsity, we also trained models on a broader set of 521 protein domains (with overall 562,208 mutations) from the Human Domainome 1 dataset. This larger dataset was used specifically for reduced-feature models containing only ESM-1v embeddings and mean DMS scores. One of the original 522 domains was excluded from this set due to its absence in MaveDB. This setup enabled investigation of model scalability and performance in scenarios where full feature availability was limited. (See Supplementary Table S1 for the full list of domains and their inclusion in the 140- and 521-domain datasets.)

All models were evaluated using RMSE, MAE, *R*^2^, and Pearson correlation (r), with group-aware train/test splitting based on gene IDs to prevent data leakage.

## 3 Results

### 3.1 Overview of Human Domainome 1 dataset

To build VEFill, we leveraged DMS data from the Human Domainome 1 dataset, which provides high-throughput mutational stability measurements across 522 protein domains. All domains were evaluated using a consistent experimental platform—an abundance-based protein fragment complementation assay (aPCA)—under standardized cellular conditions. Mutational libraries were generated using site-saturation mutagenesis, introducing all possible amino acid substitutions across each domain. After excluding one artificial sequence not present in MaveDB, 521 domains were available for downstream analysis.

From this set, we selected a subset of 140 domains (comprising 136,854 mutations) for training feature-complete models using all available data types, including EVE scores, substitution matrices, physicochemical amino acid properties, and sequence-based embeddings. This well-controlled and feature-complete subset enabled the development of VEFill as a domain-generalizable model for accurate DMS score imputation under stability-focused experimental conditions.

The full 521-domain set (comprising 562,208 mutations) was used to train reduced-feature models based solely on ESM-1v embeddings and mean DMS scores. By training reduced-feature models on this broader dataset, we explored VEFill’s performance and scalability in more practical, feature-limited scenarios.

### 3.2 General model evaluation

To investigate the influence of different feature sets on DMS score imputation, we trained a series of general models on 136,854 mutations from 140 protein domains in the Human Domainome 1 dataset using a leave-protein-out strategy, where variants from 90% of proteins were used for training and variants from the remaining 10% were held out for testing (Fig. 2, Supplementary Fig. S2, Supplementary Table S5). We evaluated performance across various biologically motivated feature combinations, including evolutionary scores (EVE), substitution matrices, physicochemical properties, and ESM-1v sequence embeddings. Simpler models using individual feature types—such as only EVE scores, substitution matrices, or biochemical descriptors—exhibited limited performance (*R*^2^ *<* 0.15). Incorporating ESM-1v embeddings led to a substantial improvement (*R*^2^ = 0.42), highlighting the utility of learned contextual sequence representations. A model relying solely on mean DMS scores per position, without any additional external features, performed better than these minimal configurations and even outperformed the model using only ESM-1v embeddings, achieving an *R*^2^ of 0.53. However, it still underperformed compared to the ESM-1v along with mean DMS model (*R*^2^ = 0.63), indicating that learned embeddings and position-specific priors capture complementary information. The best-performing model, which included all features except the ESM-1v difference vector, achieved a very modest improvement of *R*^2^ = 0.64 and a Pearson correlation of r = 0.80 (Fig. 3), suggesting that most of the relevant signal from sequence embeddings was captured by wild-type and variant ESM-1v representations alone (see Supplementary Table S3 for abbreviations used for feature sets). These findings underscore the importance of integrating biologically relevant features—particularly learned embeddings and positional priors—for accurate, cross-domain prediction of mutational effects.

**Fig. 3:**
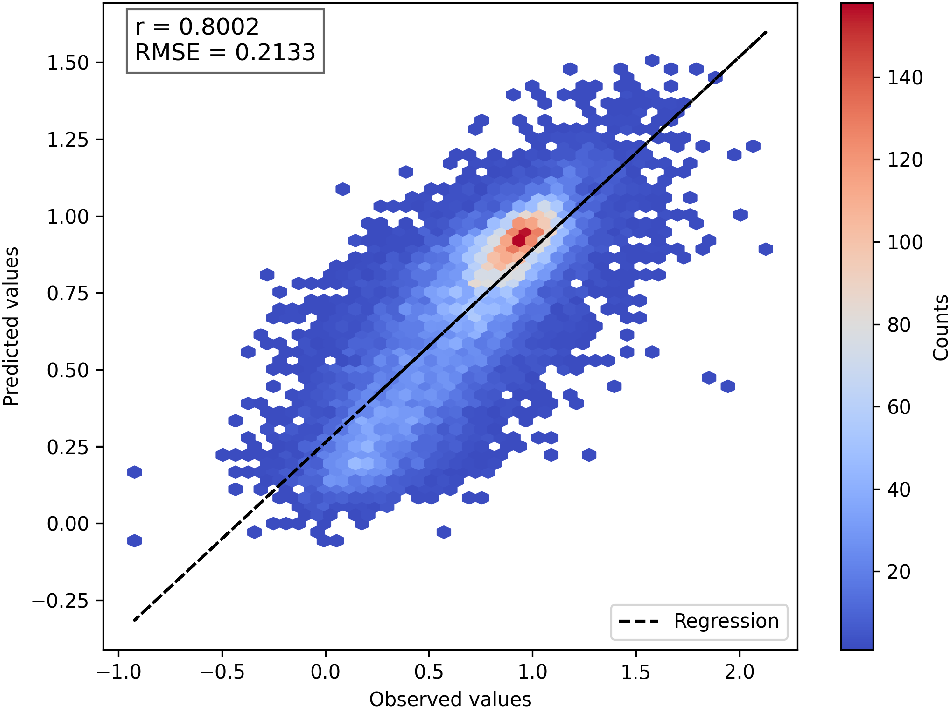
Correlation between predicted and observed DMS scores for the best general model. Predicted scores are from the model using all features except the ESM-1v difference embedding. The 2D histogram shows prediction density. Most predictions cluster around the wild-type score of 1, reflecting the prevalence of near-neutral variants. The normalized DMS scores are theoretically bounded between 0 (nonsense-like) and 1 (neutral-like), but both predicted and observed scores occasionally fall outside this range. This is due to the fact that some experimental DMS datasets contain out-of-bound values, which are preserved during normalization. The dashed line shows perfect prediction (y = x).

### 3.3 General (LOPO) model evaluation

To further evaluate generalizability, we trained 140 leave-one-protein-out (LOPO) models—each excluding one protein domain for testing. The model achieved consistently strong predictive performance across domains, with median values of *r* = 0.82, *R*^2^ = 0.66, RMSE = 0.20, and MAE = 0.15 (Fig. 4(A), Supplementary Figure S3, Supplementary Table S6)

**Fig. 4:**
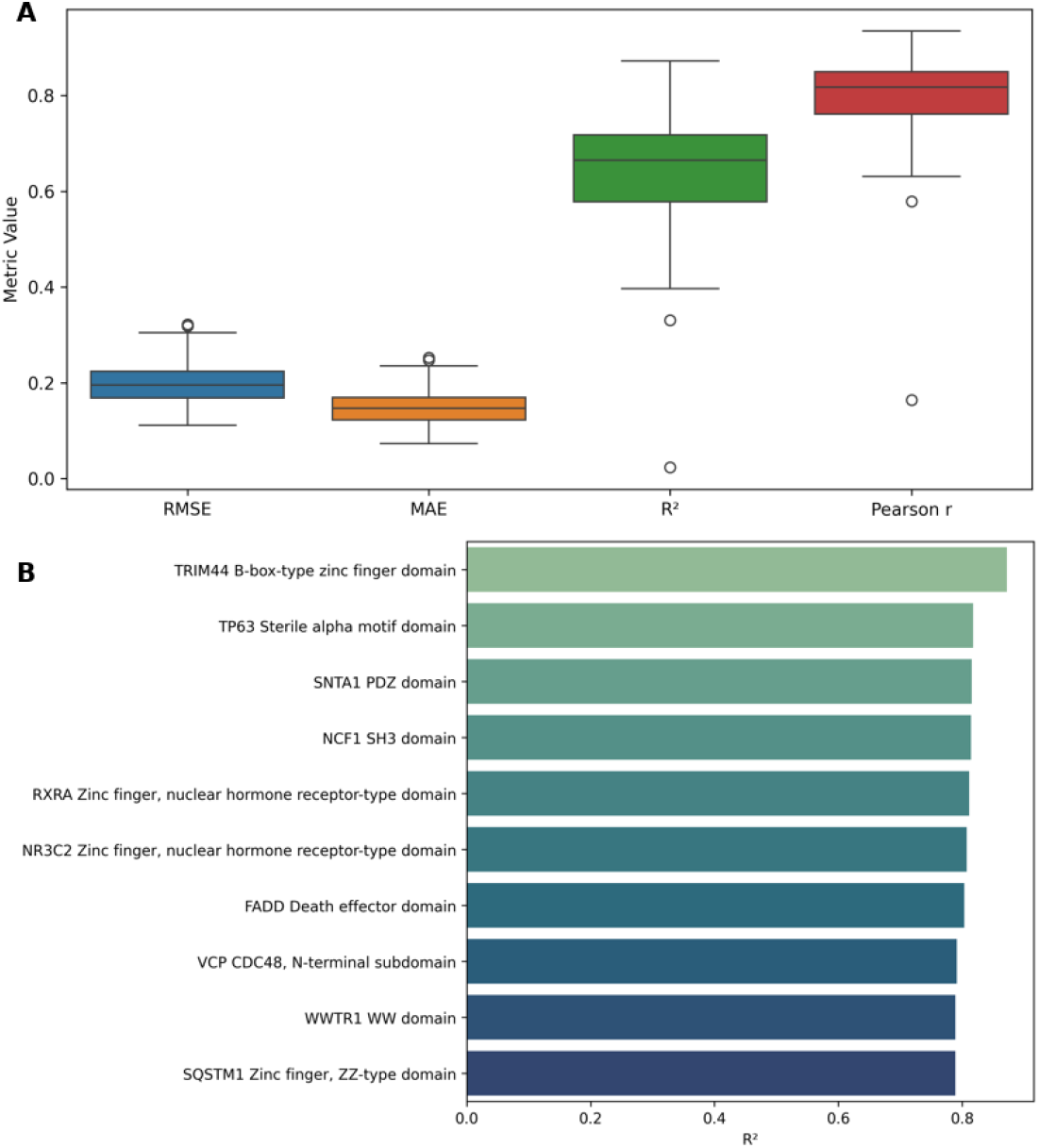
Performance evaluation of the general (LOPO) model across protein domains. (A) Evaluation metrics for the general (LOPO) model trained and tested across 140 protein domains. Each model was trained using data from 139 proteins and tested on the one held-out protein. Each box summarizes RMSE, MAE, *R*^2^, and Pearson correlation coefficient across all runs. (B) *R*^2^ values of the top 10 best-performing proteins from the same LOPO setup. Each bar represents a protein that was excluded from training and used for testing. (See Supplementary Fig. S3, Supplementary Table S6 for all individual metric values (RMSE, MAE, *R*^2^, and Pearson r) for each of the 140 LOPO models.)

To further highlight domain-specific accuracy, we identified the top 10 best-performing protein domains under the LOPO setup, based on *R*^2^ values. These domains have *R*^2^ scores between 0.78 and 0.87 (Fig. 4(B), Supplementary Figure S3, Supplementary Table S6), indicating highly reliable DMS score imputation for structurally and functionally diverse protein domains.

### 3.4 Per-protein model evaluation: random splits, LPosO, split by SNV

We evaluated the predictive performance of per-protein models across various data partitioning strategies and feature sets. As expected, models trained on a greater proportion of the data (90%, 85%, and 80%) achieved better performance (Fig. 5(A), Supplementary Fig. S4, Supplementary Tables S7, S8, S9) compared to those trained on more limited subsets (20%, 15%), underscoring the importance of data richness in training (Fig. 5(A), Supplementary Tables S10, S11). To assess robustness under data scarcity, we simulated a high-sparsity scenario by reducing training data to as little as 15%, observing a modest decline in performance. In the most extreme configuration (15% training / 85% test), the average Pearson correlation dropped by up to 0.1 compared to the richer configuration (85% training / 15% test) across the top 10 proteins (Fig. 5(A), Supplementary Tables S8, S11). These results highlight the adaptability of VEFill in low-data regimes and underscore its potential for guiding targeted mutagenesis efforts by prioritizing informative variants.

**Fig. 5:**
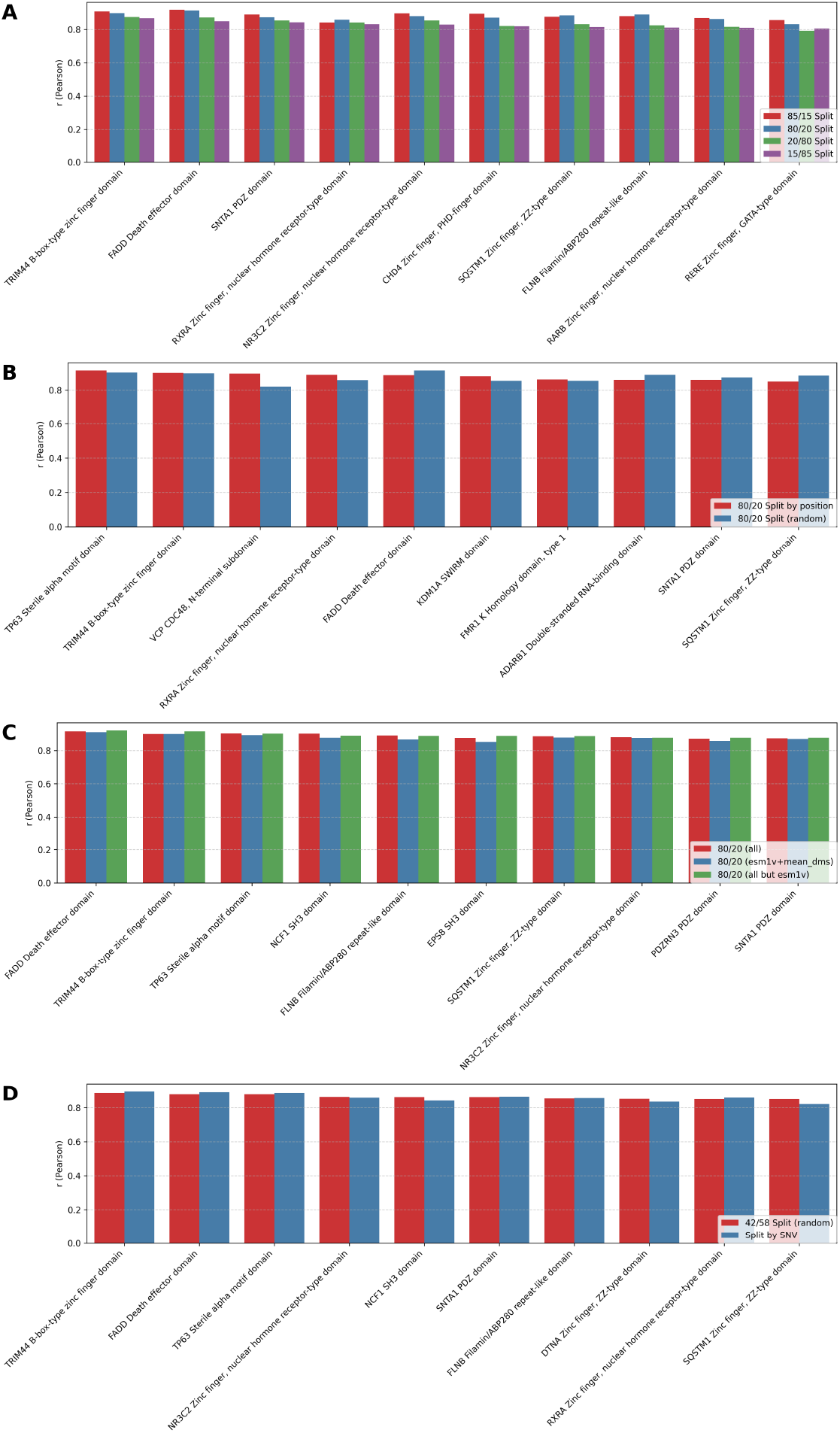
Performance of the top 10 per-protein models based on Pearson correlation coefficient (r) between observed and predicted DMS scores under various data splitting strategies and feature sets. (A) Comparison of per-protein models trained using four different random train/test splits: 85/15, 80/20, 20/80 and 15/85. Y-axis shows the r value for one protein’s model, with top 10 proteins ranked by performance in the 15/85 split. (B) Performance comparison between random 80/20 splitting and leave-position-out (LPosO) strategy for the top 10 proteins (based on LPosO performance). In LPosO, all variants at 20% of the positions are held out during training. (C) Effect of feature selection on per-protein model performance. All models use the 80/20 random split, and three feature sets are compared: **(red)** full feature set, **(blue)** ESM-1v embeddings and mean DMS score only, and **(green)** full set excluding ESM-1v. The top 10 per-protein models are selected based on their performance without ESM-1v features. (D) Comparison of model performance using two training strategies: training exclusively on SNVs, and a 42/58 random train/test split, which corresponds to the average split ratio in the SNV-based models. The top 10 models are selected based on performance with the 42/58 train/test split.

Performance comparison between the random 80/20 split and the leave-position-out (LPosO) configuration for the top 10 proteins—ranked by LPosO performance—revealed consistent trends in predictive accuracy. The LPosO setup, which simulates a more stringent scenario by withholding all variants from 20% of sequence positions during training, demonstrated the model’s capacity to generalize across distinct positions of the protein sequence. (Fig. 5(B), Supplementary Tables S9, S12). We also tested how reduced feature sets influenced model performance. Models trained using only ESM-1v and mean DMS scores, as well as models trained without ESM-1v embeddings, retained competitive performance compared to models trained on full feature set (Fig. 5(C), Supplementary Tables S9, S13, S14).

Finally, we compared performance between models trained exclusively on single-nucleotide variants (SNVs) and those trained using a 42/58 random train/test split, which reflects the average training proportion observed in the SNV-based splits. Despite training on a more restricted subset of mutations, models trained on SNVs alone achieved comparable performance to those trained on the broader mutation set, indicating that informative SNVs can capture key predictive signals (Fig. 5(D), Supplementary Tables S15, S16).

### 3.5 Fine-grained per-protein evaluation: LOPosO vs. LOVarO

To explore model generalization at finer granularity, we compared leave-one-position-out (LOPosO) and leave-one-variant-out (LOVarO) strategies on two top-performing models for FADD Death effector domain and TRIM44 B-box-type zinc finger domain. In the LOPosO setup, all variants at a single position were held out during each iteration, evaluating the model’s ability to generalize across positions. LOVarO, in contrast, held out a single variant at a time, enabling localized performance evaluation within familiar sequence contexts.

Both strategies showed consistently low error rates, affirming the model’s robustness. However, LOPosO exhibited slightly higher prediction variability, consistent with its broader generalization demands (Fig. 6).

**Fig. 6:**
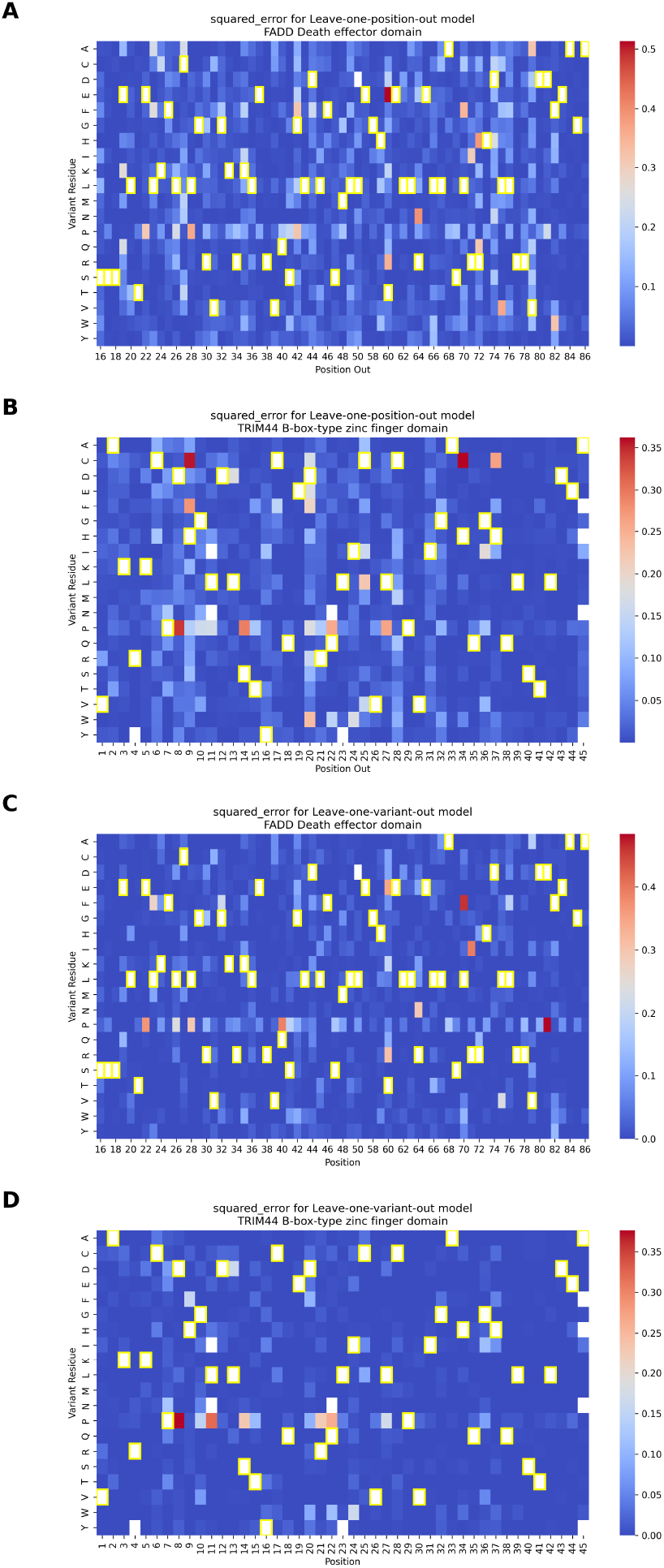
The squared error (SE) between predicted and observed normalized DMS scores for the two best-performing per-protein models—FADD Death effector domain and TRIM44 B-box-type zinc finger domain—evaluated under leave-one-position-out (LOPosO) and leave-one-variant-out (LOVarO) strategies. **(A, B)** SE distributions for FADD and TRIM44, respectively, under the LOPosO strategy, where in each iteration, all variants at a single position are held out for testing. The squared error for each variant at the held-out position is calculated and plotted in the heatmap. This process is repeated across all positions, holding out a different position in each iteration. **(C, D)** SE distributions for FADD and TRIM44, respectively, using the LOVarO strategy, in which only one variant is held out at a time. The SE is calculated for that single held-out variant and plotted accordingly. This fine-grained approach evaluates the model’s ability to predict individual unseen mutations. Cells with yellow frames indicate the wild-type amino acid at each position. (See Supplementary Tables S17-S20 for all values used for these plots along with other evaluation metrics.)

Several mutation-specific patterns emerged from the squared error distributions. Proline (P) substitutions consistently led to elevated prediction errors across both strategies, likely due to proline’s unique conformational constraints and disruption of secondary structures—effects that are not easily captured by sequence-based features such as ESM-1v embeddings.

We also observed that extremely damaging variants, particularly those with normalized scores below the loss-of-function threshold, were more prone to higher squared error. This occurred despite overall strong performance for most damaging variants, reflecting the inherent challenge of modeling rare, high-impact effects.

Interestingly, for TRIM44 B-box-type zinc finger domain, several histidine-to-cysteine (H→C) mutations were poorly predicted by LOPosO but accurately captured by LOVarO. This discrepancy highlights the value of fine-grained, position-aware evaluation in capturing context-dependent biochemical nuances. Histidine and cysteine, for instance, can coordinate zinc ions in domain-specific roles that LOPosO may misclassify due to its generalized view [33]. (Fig. 6, Supplementary Tables S17-S20, see Supplementary Fig. S5 for additional insight into correlation between predicted scores and error magnitudes, and Supplementary Note S1 for discussion on interpretation of LPOsO and LOVarO discrepancies).

### 3.6 Generalization to non-Domainome assays

The general model trained on the Human Domainome 1 dataset was further evaluated on 8 full-length, non-Domainome proteins from MaveDB that were not seen during training. The model showed substantially better performance on stability-based assays [34–36] than on activity-based ones [4, 5, 37, 38] (Table 1, Fig. 7, Supplementary Table S21, Supplementary Fig. S6-S7). This demonstrates the importance of generating experimental data for diverse assay types to enable more powerful computational models.

**Table 1.** General model performance on external assays

| Protein | Assay | RMSE | R<sup>2</sup> |
| --- | --- | --- | --- |
| PRKN [34] | stability | 0.175 | 0.799 |
| PTEN [35] | stability | 0.214 | 0.550 |
| ASPA [36] | stability | 0.229 | 0.702 |
| CALM1 [4] | activity | 0.457 | 0.152 |
| PTEN [5] | activity | 0.495 | 0.465 |
| TPK1 [4] | activity | 0.544 | 0.158 |
| TP53 [37] | activity | 0.793 | 0.473 |
| TP53 [38] | activity | 1.438 | -0.125 |
| Human Domainome 1 [18] | stability | 0.213 | 0.640 |

**Fig. 7:**
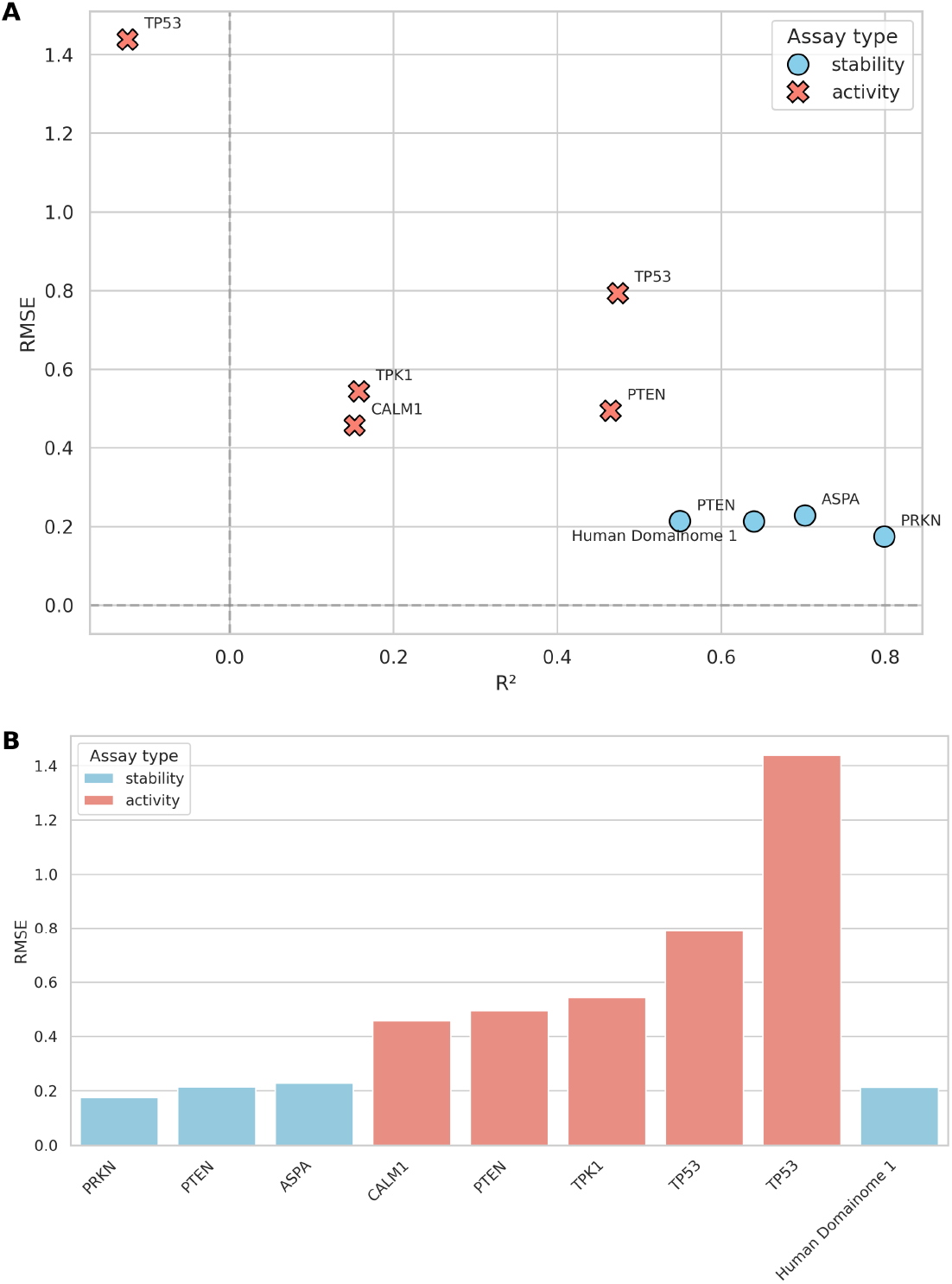
Performance of the general model across stability-based (blue) and activity-based (red) external assays. (A) compares RMSE and *R*^2^ values across assays, illustrating substantially better predictive accuracy on stability-based assays (PRKN [34], PTEN [35], ASPA [36]) compared to activity-based assays (CALM1 [4], PTEN [5], TPK1 [4], TP53 [37, 38]). (B) summarizes the differences in RMSE explicitly, highlighting the notable error increase when predicting activity-based assay data relative to stability-based data. Performance on internal validation (Human Domainome 1) data is included as a reference for comparison. (See Supplementary Fig. S6 (A–F) for additional visualizations of performance metrics (RMSE vs. Pearson r, Pearson r vs. *R*^2^, MAE vs. RMSE), and Supplementary Fig. S7 (A–H) for detailed visualizations of Pearson r, *R*^2^, and MAE, and correlations of predicted vs. observed DMS scores for each assay.)

### 3.7 Minimal feature model for imputation

We tested a lightweight version of the model that uses (i) only ESM-1v embeddings (wild-type, variant, and difference vectors) combined with mean DMS scores per position (Supplementary Table S22)—two features identified as highly important in our feature selection analysis—and (ii) mean DMS scores alone (Supplementary Table S23). Reduced-feature models were trained on either 140 or 521 protein domains from the Human Domainome 1 dataset.

We compared the performance of these models with the full-feature model using a validation set (10% holdout from Human Domainome 1) and external evaluation on full-length non-Domainome proteins (PRKN [34], multiple PTEN [5, 35], ASPA [36], CALM1 [4], TPK1 [4], and multiple TP53 [37, 38]) with 30% of mutations masked, covering both stability- and activity-based DMS datasets (Fig. 8, Supplementary Fig. S8). The full-feature model consistently outperformed the reduced versions. However, the reduced models still performed competitively, making them a practical and scalable alternative when only limited features are available.

**Fig. 8:**
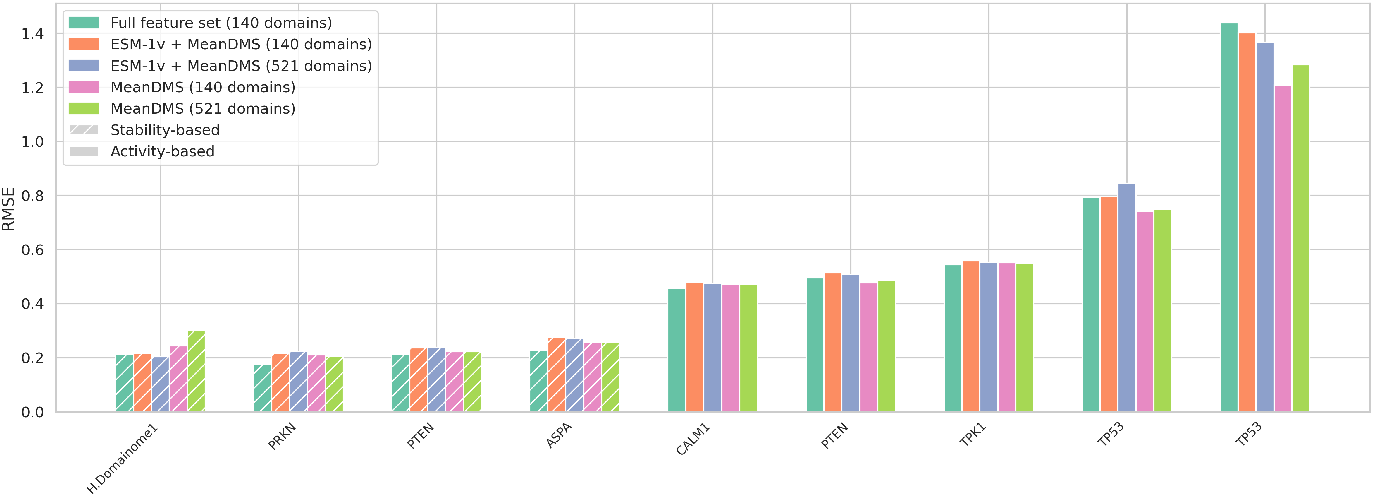
Performance comparison of full-feature and reduced-feature models using RMSE as the primary evaluation metric. The full-feature model includes all input features, while reduced models include either (i) only ESM-1v embeddings and mean DMS scores or (ii) solely mean DMS scores. All reduced-feature models were trained on either 140 or 521 protein domains from the Human Domainome 1 dataset. RMSE values are shown for internal validation on the Human Domainome 1 test set and for external full-length proteins (from left to right: PRKN [34], PTEN [35], ASPA [36], CALM1 [4], PTEN [5], TPK1 [4], TP53 [37, 38]) assayed across multiple platforms. Patterned bars denote stability-based assays, which consistently yield stronger predictive performance. The full-feature model outperforms the reduced models across nearly all datasets.

On the internal test set, the ESM-1v + mean DMS model consistently outperformed the mean DMS-only version. However, this advantage did not consistently translate to the full-length protein dataset, where both reduced models performed comparably across most non-Domainome proteins. Similarly, increasing the training set size from 140 to 521 domains did not yield substantial changes in performance.

To better understand these results, we analyzed the representation of protein families across the training and test sets of the 140- and 521-domain datasets using the Pfam database [39] (Fig. 9). We observed a skewed distribution where a few dominant Pfam families (e.g., PF00046, PF00018) contributed a large fraction of total protein domains. In the 521-domain dataset, the top 5 Pfam families alone accounted for over 30% of all protein domains. Such overrepresentation introduces redundancy, potentially explaining why increasing the training size from 140 to 521 domains did not yield major improvements in external generalization.

**Fig. 9:**
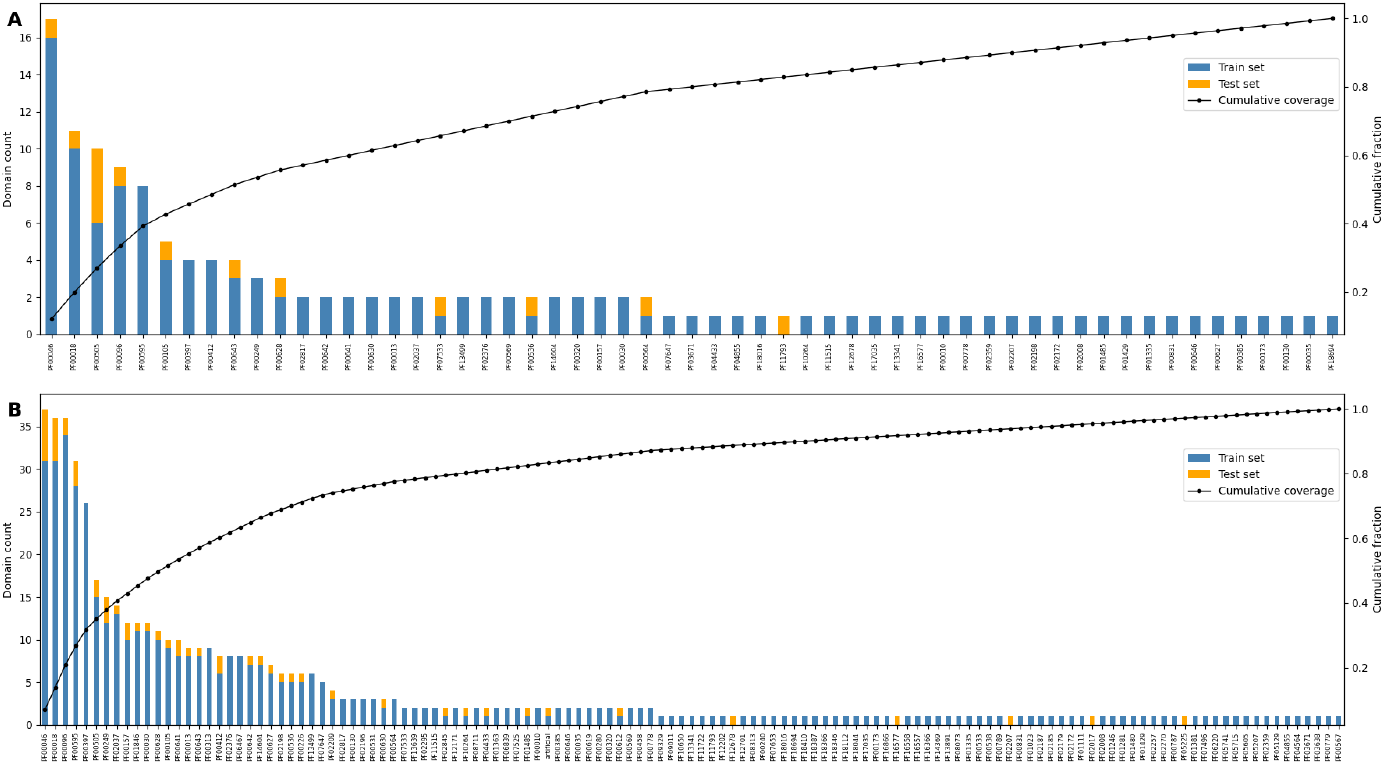
Distribution of protein families in the 140-domain and 521-domain training datasets used for reduced-feature model training. **(A)** 140-domain dataset; **(B)** 521-domain dataset. The left Y-axis indicates the number of protein domains per protein family. Bars are stacked to show counts in the training (blue) and test (orange) sets. Pfam IDs are ordered by descending total domain count. The right Y-axis shows the cumulative fraction of the dataset accounted for by each Pfam family (black line), highlighting the dominance of a small subset of Pfam families. (See Supplementary Table S1 for detailed list of domains, including UniProt IDs, Pfam IDs, and their presence in the 140- and 521-domain datasets.)

To directly test the effect of this redundancy, we constructed a reduced dataset from Human Domainome 1 containing one protein domain per Pfam family (127 domains in total, see Supplementary Table S1 for the full list of domains and their inclusion in the 140-, 521-, and 127-domain (Pfam-unique) datasets), ensuring complete non-overlap of Pfam families between training and test sets. Models trained on this Pfam-unique set exhibited slightly higher RMSE compared to those trained on the full 521-domain set: 0.2654 vs. 0.2039 for the ESM-1v + mean DMS model, and 0.3671 vs. 0.3012 for the mean DMS-only model. These modest increases suggest that while Pfam redundancy exists, it does not strongly bias performance or lead to overfitting in our reduced-feature models.

### 3.8 Zero-shot generalization (no mean DMS)

To assess the model’s potential for zero-shot prediction, we evaluated a general model trained without the positional mean DMS score feature on external, non-Domainome datasets. Since the mean DMS score per position is derived directly from experimental data, removing this feature ensures no data leakage from training to testing—a necessary condition for genuine zero-shot evaluation.

We tested the model on both stability-based (PRKN, PTEN, ASPA) and activity-based (CALM1, PTEN, TPK1, TP53) external assays (Supplementary Table S24). While performance decreased compared to the full-feature version (Table 1, Fig. 7, Supplementary Table S21, Supplementary Fig. S6-S7), especially on activity-based assays, the model still demonstrated relatively better accuracy on stability-based over activity-based datasets (Fig. 10, Supplementary Fig. S9-S10). These results confirm that although the model can still capture useful general patterns, its accuracy without access to local experimental context (mean DMS scores) remains limited.

**Fig. 10:**
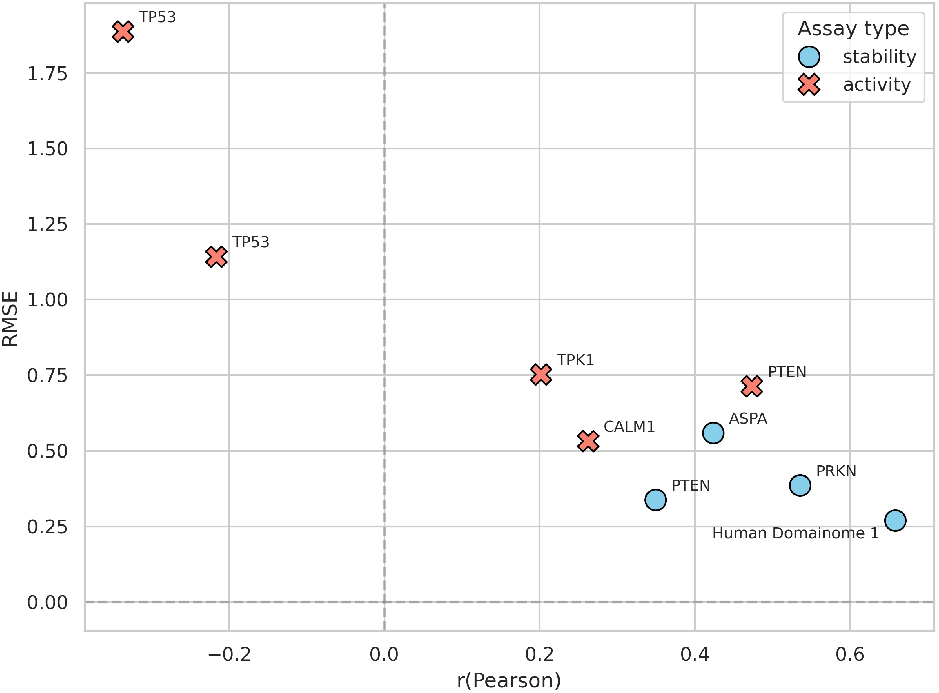
Zero-shot general model (without mean DMS) performance: RMSE vs. Pearson correlation (r) of zero-shot predictions by the general model trained without positional mean DMS scores. Stability-based assays (blue circles) consistently exhibit better predictive accuracy than activity-based assays (red crosses). The internal Human Domainome 1 validation is shown for reference. (See Supplementary Fig. S9 (A–C) for further details on additional metrics (RMSE vs. *R*^2^, Pearson r vs. *R*^2^, MAE vs. RMSE), and Supplementary Fig. S10 (A–H) for correlations of predicted vs. observed DMS scores for each assay.)

## 4 Discussion

Our findings demonstrate that VEFill—a LightGBM-based model integrating evolutionary, biochemical, and sequence-based features—enables accurate imputation of deep mutational scanning (DMS) scores across a diverse range of human protein domains. Trained on the standardized and stability-focused Human Domainome 1 dataset, VEFill achieved strong predictive performance, with the highest gains observed when using contextual embeddings (ESM-1v) and position-specific priors (mean DMS score per position). These results underscore the utility of combining data-driven positional information and learned sequence representations to enhance model generalization.

Evaluation on both internal and external datasets confirmed VEFill’s capacity to generalize across previously unseen domains assayed under stability-focused experimental conditions. However, its performance on activity-based datasets was markedly reduced, likely due to mismatches between training and test assay modalities, as well as the complex nature of cellular phenotypes.

Performance assessments of VEFill under various per-protein train–test configurations (random splits, LPosO, LOPosO, LOVarO) showed strong robustness in sparse-data scenarios. LOVarO consistently outperformed other strategies, particularly for context-specific or chemically subtle mutations, underscoring the importance of maintaining positional information during training. Notably, substitutions involving proline residues posed consistent prediction challenges, likely due to their disruptive conformational effects not fully captured by the feature set. Additionally, extremely deleterious variants—especially those falling below the normalized loss-of-function threshold—tended to have higher prediction error, despite overall strong model performance on damaging mutations. This reflects the inherent difficulty of modeling rare, high-impact outliers. Specific mutation types, such as histidine-to-cysteine substitutions in zinc finger domains, were predicted more accurately under fine-grained LOVarO validation compared to LOPosO, further emphasizing the importance of preserving positional and structural nuances (see Supplementary Note S1).

A reduced version of VEFill, incorporating only ESM-1v embeddings and mean DMS scores, achieved performance comparable to the full model in most cases. This lightweight configuration may be particularly useful in resource-constrained settings. However, excluding position-level priors significantly reduced accuracy in zero-shot scenarios, highlighting their critical role in enabling generalizable variant effect prediction.

To further investigate model generalization, we evaluated the performance of reduced-feature models trained on either 140 or 521 protein domains and analyzed the distribution of Pfam families across these training datasets. A highly skewed distribution was observed, where a small number of Pfam families dominated the dataset. In the 521-domain set, for instance, the top five Pfam families alone accounted for over 30% of all protein domains. This biological redundancy reflects true enrichment of certain protein families in the human genome, but it also limits family diversity. As a result, increasing the training size from 140 to 521 domains mainly added examples from already well-represented families, offering limited coverage of novel Pfam families. Consequently, the larger dataset did not confer clear improvements in external generalization, particularly when test proteins belonged to families absent from the training data.

We emphasize that protein family overlap between training and test sets is not a data leakage artifact but a biologically relevant feature of protein space. Domains within the same Pfam family tend to share structural and functional properties, and generalizing within families is desirable. To explicitly test whether shared Pfam families may inflate performance, we repeated training using only one protein domain per Pfam family (127 total), thereby eliminating any Pfam-level leackage between train and test splits. RMSE increased only modestly (by 0.07 for both reduced-feature models), suggesting that while redundancy contributes to model learning, it is not a potential source of overfitting. These results support the robustness of our findings and reinforce that increasing Pfam diversity, rather than dataset size alone, may be more important for generalization across structurally unrelated proteins.

VEFill offers a scalable and interpretable framework for DMS score imputation. Its effectiveness in low-data regimes suggests utility in guiding experimental design, prioritizing mutations for targeted assays, and constructing efficient mutational libraries. Beyond current capabilities, the model could be further improved by explicitly incorporating structural features. Although VEFill does not directly use protein structure, ESM-1v embeddings may implicitly encode structural patterns. Nonetheless, we observed a decline in predictive performance at structurally sensitive positions—such as proline substitutions—indicating that the explicit inclusion of structural features may further improve model accuracy.

Future directions include incorporating structural modeling predictions or annotations to address context-specific error patterns. Cross-species transfer learning, leveraging multiple sequence alignments, may also extend VEFill’s utility to predict variant effects in non-human proteins using experimental data from related species.

## 5 Availability of source code and requirements

**Project name:** VEFill

**Project homepage:** https://github.com/PlushZ/VEFill

**Source code:** available at Zenodo [40]

**Operating system:** Platform independent

**Programming language:** Python

**Other requirements:** Python 3.10 or higher

**License:** MIT license

## 6 Data Availability

All data used in this study are available at Zenodo [41]:

- `db/` contains a PostgreSQL backup file. It can be restored using `pg_restore` and includes all tables used to train VEFill.
- `data/` contains processed datasets used in this study, including both Domainome and non-Domainome data. These datasets contain mutation-level features and target labels used to train and evaluate the machine learning models described in the manuscript. The data was generated using the preprocessing pipeline available in the accompanying code repository.
- `models/` contains the pretrained VEFill models.
- `supplementary/` contains all supplementary materials for this manuscript.

Each component can be used and reproduced with the accompanying code.

## 7 Additional Files

**Supplementary Table S1**. Metadata for domains used in the 140-, 521-, and 127-domain (Pfam-unique) model datasets. Includes MaveDB accession, domain name, species, UniProt ID, Pfam ID. Train/test split assignments are indicated for each dataset where applicable.

**Supplementary Table S2**. List of the 25 amino acid substitution matrices used in VEFill. Each row includes the matrix name, its source (Biopython or external repository), the corresponding VEFill feature name, and the original citation.

**Supplementary Table S3**. Complete list of input features used in the VEFill model.

**Supplementary Table S4**. Performance comparison of different model architectures using the same input feature set. Models evaluated include LightGBM (used in VEFill), a fully connected neural network (FCNN), and a transformer-based architecture. Reported metrics include RMSE, MAE, and *R*^2^ for both training and test sets. Variants of the FCNN included adjustments to regularization, architecture depth, and optimizer settings. All models were trained using the same 140-domain dataset.

**Supplementary Table S5**. Performance metrics (RMSE, MAE, *R*^2^, Pearson r) for both the training and test sets of the general VEFill model, evaluated across multiple feature set configurations. Each row represents a unique feature combination used to evaluate model performance.

**Supplementary Table S6**. Domain-wise evaluation results for the general model under a LOPO strategy. The table reports RMSE, MAE, coefficient of determination (*R*^2^), and Pearson correlation (r) for both the training and test sets across 140 individual protein domains. Additionally, the number of mutations in the training and test sets for each domain is provided.

**Supplementary Table S7**. Per-protein model performance metrics (RMSE, MAE, *R*^2^, Pearson r) for both training and test sets for 140 domains using an 90/10 random train/test split. Models were trained using all input features except ESM-1v difference vector. The table also includes the number of mutations in the training and test sets for each domain.

**Supplementary Table S8**. Per-protein model performance metrics (RMSE, MAE, *R*^2^, Pearson r) for both training and test sets for 140 domains using an 85/15 random train/test split. Models were trained using all input features except ESM-1v difference vector. The table also includes the number of mutations in the training and test sets for each domain.

**Supplementary Table S9**. Per-protein model performance metrics (RMSE, MAE, *R*^2^, Pearson r) for both training and test sets for 140 domains using an 80/20 random train/test split. Models were trained using all input features except ESM-1v difference vector. The table also includes the number of mutations in the training and test sets for each domain.

**Supplementary Table S10**. Per-protein model performance metrics (RMSE, MAE, *R*^2^, Pearson r) for both training and test sets for 140 domains using an 20/80 random train/test split. Models were trained using all input features. The table also includes the number of mutations in the training and test sets for each domain.

**Supplementary Table S11**. Per-protein model performance metrics (RMSE, MAE, *R*^2^, Pearson r) for both training and test sets for 140 domains using an 15/85 random train/test split. Models were trained using all input features except ESM-1v difference vector. The table also includes the number of mutations in the training and test sets for each domain.

**Supplementary Table S12**. Per-protein model performance metrics (RMSE, MAE, *R*^2^, Pearson r) for both training and test sets across 140 protein domains, using an 80/20 train/test split under the LPosO strategy. Models were trained using all input features. The table also reports the number of amino acid positions and mutations in the training and test sets for each domain.

**Supplementary Table S13**. Per-protein model performance metrics (RMSE, MAE, *R*^2^, Pearson r) for both training and test sets across 140 protein domains, using an 80/20 random train/test split. Models were trained using only ESM-1v embeddings and mean DMS score as input features. The table also includes the number of mutations in the training and test sets for each domain.

**Supplementary Table S14**. Per-protein model performance metrics (RMSE, MAE, *R*^2^, Pearson r) for both training and test sets across 140 protein domains, using an 80/20 random train/test split. Models were trained using all input features except ESM-1v embeddings. The table also includes the number of mutations in the training and test sets for each domain.

**Supplementary Table S15**. Per-protein model performance metrics (RMSE, MAE, *R*^2^, Pearson r) for both training and test sets across 140 domains using a split based on the SNV indicator. The training set includes only data points with an SNV indicator of 1, while the test set includes those with an SNV indicator of 0. Models were trained using all input features except the SNV indicator. The table also reports the number of mutations in the training and test sets for each domain.

**Supplementary Table S16**. Per-protein model performance metrics (RMSE, MAE, *R*^2^, Pearson r) for both training and test sets for 140 domains using an 42/58 random train/test split. Models were trained using all input features. The table also includes the number of mutations in the training and test sets for each domain.

**Supplementary Table S17**. Prediction error metrics for the FADD Death effector domain using the LOPosO strategy. For each held-out position, the table reports observed and predicted DMS scores, absolute error, squared error, percentage error, and the corresponding wild-type and variant residues.

**Supplementary Table S18**. Prediction error metrics for the TRIM44 B-box-type zinc finger domain using the LOPosO strategy. For each held-out position, the table reports observed and predicted DMS scores, absolute error, squared error, percentage error, and the corresponding wild-type and variant residues.

**Supplementary Table S19**. Prediction error metrics for the FADD Death effector domain using the LOVarO strategy. Each row corresponds to one held-out variant and includes observed and predicted DMS scores, absolute error, squared error, percentage error, position, wild-type residue, and variant residue.

**Supplementary Table S20**. Prediction error metrics for the TRIM44 B-box-type zinc finger domain using the LOVarO strategy. Each row corresponds to one held-out variant and includes observed and predicted DMS scores, absolute error, squared error, percentage error, position, wild-type residue, and variant residue.

**Supplementary Table S21**. Performance metrics of the general model (trained on the Human Domainome 1 dataset) tested on external stability-based (PRKN, PTEN, ASPA) and activity-based (CALM1, PTEN, TPK1, TP53) DMS assays. Each dataset was evaluated by masking 30% of the experimental DMS scores and predicting their values. Metrics shown include RMSE, MAE, *R*^2^, and Pearson r. The final row shows the performance metrics of the general model on Human Domainome 1 internal validation data, serving as a benchmark for comparison.

**Supplementary Table S22**. Performance metrics for the general 140- and 521-domain models trained on a reduced feature set (including only mean DMS scores and ESM-1v embeddings), tested on external stability-based (PRKN, PTEN, ASPA) and activity-based (CALM1, PTEN, TPK1, TP53) assays. Metrics include RMSE, MAE, *R*^2^, Pearson r. Performance on the internal Human Domainome 1 test set using the same reduced feature set is included for comparison.

**Supplementary Table S23**. Performance metrics for the general 140- and 521-domain models trained on a reduced feature set (including only mean DMS scores), tested on external stability-based (PRKN, PTEN, ASPA) and activity-based (CALM1, PTEN, TPK1, TP53) assays. Metrics include RMSE, MAE, *R*^2^, Pearson r. Performance on the internal Human Domainome 1 test set using the same reduced feature set is included for comparison.

**Supplementary Table S24**. Performance metrics for the zero-shot general model trained without positional mean DMS score, tested on external stability-based (PRKN, PTEN, ASPA) and activity-based (CALM1, PTEN, TPK1, TP53) assays. Metrics include RMSE, MAE, *R*^2^, Pearson r. The final row represents performance metrics from internal validation on the Human Domainome 1 dataset (also without mean DMS scores), provided for comparison.

**Supplementary Fig. S1**. Schema of the custom PostgreSQL database developed for centralized storage and structured access to all model input features.

**Supplementary Fig. S2**. Performance metrics of the general model trained and evaluated on various feature sets for DMS score imputation. Metrics shown include RMSE, MAE, R-squared (*R*^2^), and Pearson correlation coefficient (r). Feature sets on the x-axis are sorted by increasing Pearson correlation coefficient, highlighting the progressive improvements achieved by enriching the model with additional informative features. Abbreviations used in figure: esm: All ESM-1v embeddings (wild-type, variant, difference); wt esm: ESM-1v embeddings for wild-type amino acid; var esm: ESM-1v embeddings for variant amino acid; diff esm: Difference between ESM-1v embeddings for wild-type and variant amino acid; mean dms: Mean DMS score per amino acid position; jse dms: James-Stein estimator of DMS score per position (global mean calculated per protein); pc: Physico-chemical properties (wild-type, variant, and their difference); sm: All substitution matrices (25 matrices); top5sm: 5 substitution matrices (‘blosum62’, ‘blosum80’, ‘blosum90’, ‘grantham’, ‘gonnet1992’); snv: Single-nucleotide variant indicator; eve: EVE scores.

**Supplementary Fig. S3**. Heatmap showing the performance of the general model across 140 individual protein domains, each evaluated using a leave-one-protein-out (LOPO) strategy. For each protein, the model was trained on all other domains and tested on the held-out domain. Shown metrics include RMSE, MAE, *R*^2^, Pearson r. Each row represents a single protein domain, with separate columns for training and test performance metrics. Color encodes the metric value: warmer colors indicate higher values—reflecting better performance for *R*^2^ and r, but poorer performance for RMSE and MAE—while cooler colors indicate lower values—meaning better performance for RMSE and MAE, and worse performance for *R*^2^ and Pearson r. **Supplementary Fig. S4**. Per-protein model performance and dataset sizes across protein domains. Heatmap on the left displays the training and testing performance of per-protein models trained with an 80/20 train/test split across 140 protein domains. Each row represents a single protein domain. Metrics shown include RMSE, MAE, *R*^2^, and Pearson r, with separate columns for training and test results. Warmer shades indicate higher values—desirable for *R*^2^ and r, but undesirable for RMSE and MAE—while cooler shades indicate lower values. Accompanying heatmap on the right illustrates the corresponding train and test set sizes (number of mutations per protein), with brighter colors reflecting larger datasets. Together, these plots help visualize the relationship between dataset size and model performance across diverse protein domains.

**Supplementary Fig. S5**. Correlation between average observed DMS scores and average prediction error across amino acid positions. **(A–B)** Leave-one-position-out (LOPosO) model: Each point represents a single amino acid position. The x-axis shows the mean normalized DMS score (y true mean) for all variants at that position, and the y-axis shows the mean squared error (squared error mean) of model predictions for those variants. **(C–D)** Leave-one-variant-out (LOVarO) model: Each point again represents a single position, aggregating prediction errors across variants at that position where each variant was excluded once during training. Strong negative correlations (Pearson r = -0.60 and -0.67) in LOPosO models for FADD Death effector domain **(A)** and TRIM44 B-box-type zinc finger domain **(B)**, respectively, indicate that positions with lower (more damaging) DMS scores tend to have higher prediction errors. This suggests that LOPosO models struggle significantly when predicting mutations at structurally or functionally critical positions due to the lack of positional context during training. Weaker negative correlations (Pearson r = -0.35 and -0.16) in LOVarO models for the same proteins, FADD **(C)** and TRIM44 **(D)**, demonstrate improved prediction accuracy when positional context is retained. The reduced error in these models highlights their enhanced capability to capture position-specific stability effects.

**Supplementary Fig. S6. (A–F)** comparing predictive performance metrics of the general model (trained on the Human Domainome 1 dataset) across external stability-based, (blue circles/bars) and activity-based assays (red crosses/bars). Metrics compared include RMSE, Pearson r, *R*^2^, and MAE. **(A)** RMSE versus Pearson correlation (r). **(B)** Pearson correlation (r) versus *R*^2^. **(C)** MAE versus RMSE. **(D–F)** showing Pearson correlation (r) **(D)**, *R*^2^ **(E)**, and MAE **(F)** individually for each assay.

**Supplementary Fig. S7. (A–H)** comparing predicted and experimentally measured (true) normalized DMS scores for the general model trained on the Human Domainome 1 dataset, tested on external stability-based assays **(A–C)** and activity-based assays **(D–H)**. Each plot represents the density of observations by color intensity, with the black dashed line indicating perfect predictions (*y* = *x*). Stability-based assays **(A–C)** show strong predictive accuracy, with high correlation values (Pearson *r* ≥ 0.74) and low RMSE, indicating robust generalization to unseen stability data: **(A)** PTEN, **(B)** PRKN, **(C)** ASPA. Activity-based assays **(D–H)** generally exhibit weaker predictive performance, with higher RMSE and variability in correlation: **(D)** TP53: High density near loss-of-function (∼ 0) variants indicates good loss-of-function predictions, but poor prediction accuracy for functional mutations (scores between 0.5–1) and gain-of-function variants (scores *>*1, predicted close to wild-type, but clinically damaging). **(E)** CALM1: Low correlation and high prediction error indicate difficulty in predicting functional assay results. **(F)** TPK1: Low predictive accuracy and correlation highlight limitations in generalizing to functional data. **(G)** PTEN (activity-based assay): Moderate correlation but higher RMSE compared to its stability counterpart **(A)**, emphasizing assay-dependent differences in predictive accuracy. **(H)** TP53: Large RMSE and broad dispersion underscore poor prediction performance, particularly for gain-of-function mutations.

**Supplementary Fig. S8**. Performance of full-feature set vs. reduced-feature set 140- and 521-domain models on Human Domainome 1 and unseen full-length proteins. Subfigures **A-C** present the evaluation metrics used to compare model performance across both held-out Human Domainome 1 domains and full-length non-Domainome proteins: PRKN, PTEN, ASPA, CALM1, TPK1, and TP53. Bars show results for the full-feature model, ESM-1v + mean DMS model, and mean DMS-only models trained on 140 or 521 domains. Stability-based assays are visually distinguished using patterned bar fills. Subfigures: **(A)** MAE **(B)** Coefficient of Determination (*R*^2^) **(C)** Pearson’s correlation coefficient (r).

**Supplementary Fig. S9**. Performance metrics for the zero-shot general model trained without positional mean DMS scores, evaluated on external stability-based (blue circles) and activity-based (red crosses) assays. **(A)** RMSE versus *R*^2^. **(B)** Pearson correlation (r) versus *R*^2^. **(C)** MAE versus RMSE. Across all metrics, stability-based assays generally exhibit higher predictive accuracy (lower RMSE and MAE, higher Pearson correlations and *R*^2^) compared to activity-based assays. Internal validation on Domainome dataset is provided as a baseline reference.

**Supplementary Fig. S10. (A–H)** showing zero-shot predictions (without using positional mean DMS scores) versus experimentally measured (true) normalized DMS scores for stability-based (A–C) and activity-based (D–H) external assays. Color intensity indicates data density, and dashed lines represent perfect predictions (y = x). Stability-based assays **(A–C)**: **(A)** PTEN, **(B)** PRKN, **(C)** ASPA, showing moderate predictive correlations (Pearson r ∼ 0.34 to 0.54) but reduced accuracy compared to the full-featured general model (see Supplementary Fig. S7). Activity-based assays **(D–H)**: **(D)** TP53, **(E)** CALM1, **(F)** TPK1, **(G)** PTEN function assay, and **(H)** TP53, illustrating significantly weaker performance, higher prediction errors, and very low or negative correlations.

**Supplementary Note S1**. Interpretation of LOPosO and LOVarO error distributions and discrepancies between model predictions and experimental DMS scores.

## List of Abbreviations

aPCA: Abundance-based Protein Complementation Assay
DMS: Deep Mutational Scanning
ESM-1v: Evolutionary Scale Modeling (protein language model)
EVE: Evolutionary Model of Variant Effect
LightGBM: Light Gradient Boosting Machine
LOPO: Leave-One-Protein-Out
LOPosO: Leave-One-Position-Out
LOVarO: Leave-One-Variant-Out
LPosO: Leave-Position-Out
MAE: Mean Absolute Error
MAVE: Multiplexed Assay of Variant Effect
MaveDB: Multiplexed Assay of Variant Effect Database
Optuna: Bayesian Hyperparameter Optimization Framework
Pfam: Protein Families Database
r: Pearson Correlation Coefficient
R^2^: Coefficient of Determination
RMSE: Root Mean Squared Error
SNV: Single Nucleotide Variant
TPE: Tree-structured Parzen Estimator
VE: Variant Effect
VEFill: Variant Effect Fill – the imputation model presented in this study
WT: Wild-Type

## 9 Competing interests

The authors declare that they have no competing interests.

## 10 Funding

PVP and WM were supported by the Freiburg Galaxy Team funded by the German Federal Ministry of Education and Research BMBF grant 031 A538A de.NBI-RBC and the Ministry of Science, Research and the Arts Baden-Württemberg (MWK) within the framework of LIBIS/de.NBI Freiburg. AFR received funding from NIH/NHGRI grants UM1HG011969 and RM1HG010461. This project received grant funding from the Australian government.

## 11 Authors’ contributions

PVP, WM, and AFR conceived the project; PVP developed the software; PVP analyzed the data; PVP, WM, and AFR participated in data interpretation; PVP, WM, and AFR wrote the manuscript; WM and AFR supervised the project. All authors reviewed and approved the manuscript.

## 12 Acknowledgements

The computing infrastructure was partly provided by the de.NBI Cloud within the German Network for Bioinformatics Infrastructure (de.NBI) and ELIXIR-DE (Forschungszentrum Jülich and W-de.NBI-001, W-de.NBI-004, W-de.NBI-008, W-de.NBI-010, W-de.NBI-013, W-de.NBI-014, W-de.NBI-016, W-de.NBI-022).

## Notes

https://doi.org/10.5281/zenodo.16877556

https://doi.org/10.5281/zenodo.15408793

https://github.com/PlushZ/VEFill/

### Competing Interest Statement

The authors have declared no competing interest.

### Summary of Updates

Updated text, revised figures and tables, expanded supplementary material, and additional analyses to support conclusions.

