## Supplementary material for "VEFill: a model for accurate and generalizable deep mutational scanning score imputation across protein domains": Supplementary_Note_S1.pdf

### Interpretation of LOPosO and LOVarO error distributions and model discrepancies

As shown in Fig. 6, across both LOPosO (Leave-One-Position-Out) and LOVarO (Leave-One-Variant-Out) evaluations, proline substitutions consistently led to elevated prediction errors compared to other amino acids, regardless of their functional impact. This trend suggests that proline variants are intrinsically harder to model, likely due to their unique biophysical properties. Proline's rigid cyclic backbone introduces conformational constraints and often disrupts secondary structures—effects that may not be well captured by ESM-1v embeddings, which are based on evolutionary sequence patterns rather than structural dynamics.

Examining the worst-performing predictions from the heatmaps (Fig. 6) reveals another notable pattern: extremely damaging variants, especially those with DMS scores falling below the normalized loss-of-function threshold, often exhibit higher squared error, despite most damaging mutations still being reasonably well predicted. Selected examples from the FADD Death effector domain illustrate this (*Note*: DMS scores in original scale [-1,0], where -1 is nonsense score, 0 is WT score):

- T60E (DMS: -1.02): SE = 0.51 (LOPosO), 0.35 (LOVarO)
- E22P (DMS: -1.07): SE = 0.31 (LOPosO), 0.37 (LOVarO)
- D81P (DMS: -1.00): SE = 0.23 (LOPosO), 0.48 (LOVarO)
- L70F (DMS: -1.02): SE = 0.35 (LOPosO), 0.46 (LOVarO)
- R64N (DMS: -0.90): SE = 0.39 (LOPosO), 0.29 (LOVarO)

For the TRIM46 B-box-type zinc finger domain:

- D8P (DMS: -0.83): SE = 0.34 (LOPosO), 0.38 (LOVarO)

Interestingly, several histidine-to-cysteine (H→C) mutations are poorly predicted by LOPosO but accurately predicted by LOVarO, reflecting the benefits of fine-grained, position-aware validation:

- H34C (DMS: -0.10): SE = 0.36 (LOPosO), 0.015 (LOVarO)
- H9C (DMS: -0.06): SE = 0.36 (LOPosO), 0.02 (LOVarO)

- H37C (DMS: -0.10): SE = 0.26 (LOPosO), 0.03 (LOVarO)

This discrepancy emphasizes that local structural context often overrides global mutation trends. In zinc finger domains, histidine and cysteine can both coordinate $Zn^{2+}$ and act as interchangeable ligands. The LOPosO model, which generalizes over entire positions, may misclassify such substitutions due to a lack of positional nuance, whereas LOVarO, which evaluates individual variants, better captures these subtleties.

These mismatches between predicted and experimental scores do not necessarily reflect model failure—they may instead reveal biologically interesting edge cases, such as:

- Metal-binding residues with flexible ligand capacity (e.g., His↔Cys)
- Structural "hotspots" where even conservative mutations have exaggerated effects
- Gain-of-function or stabilizing mutations with out-of-scale DMS scores not well represented in the training set
- Regions under dynamic structural constraints not encoded in sequence-only features

Rather than being dismissed as noise, these outliers suggest opportunities for discovering new mechanistic insights and improving future structure- or dynamics-aware models.
