## Supplementary figures and images for "VEFill: a model for accurate and generalizable deep mutational scanning score imputation across protein domains"

### Supplementary_Fig_S1.png

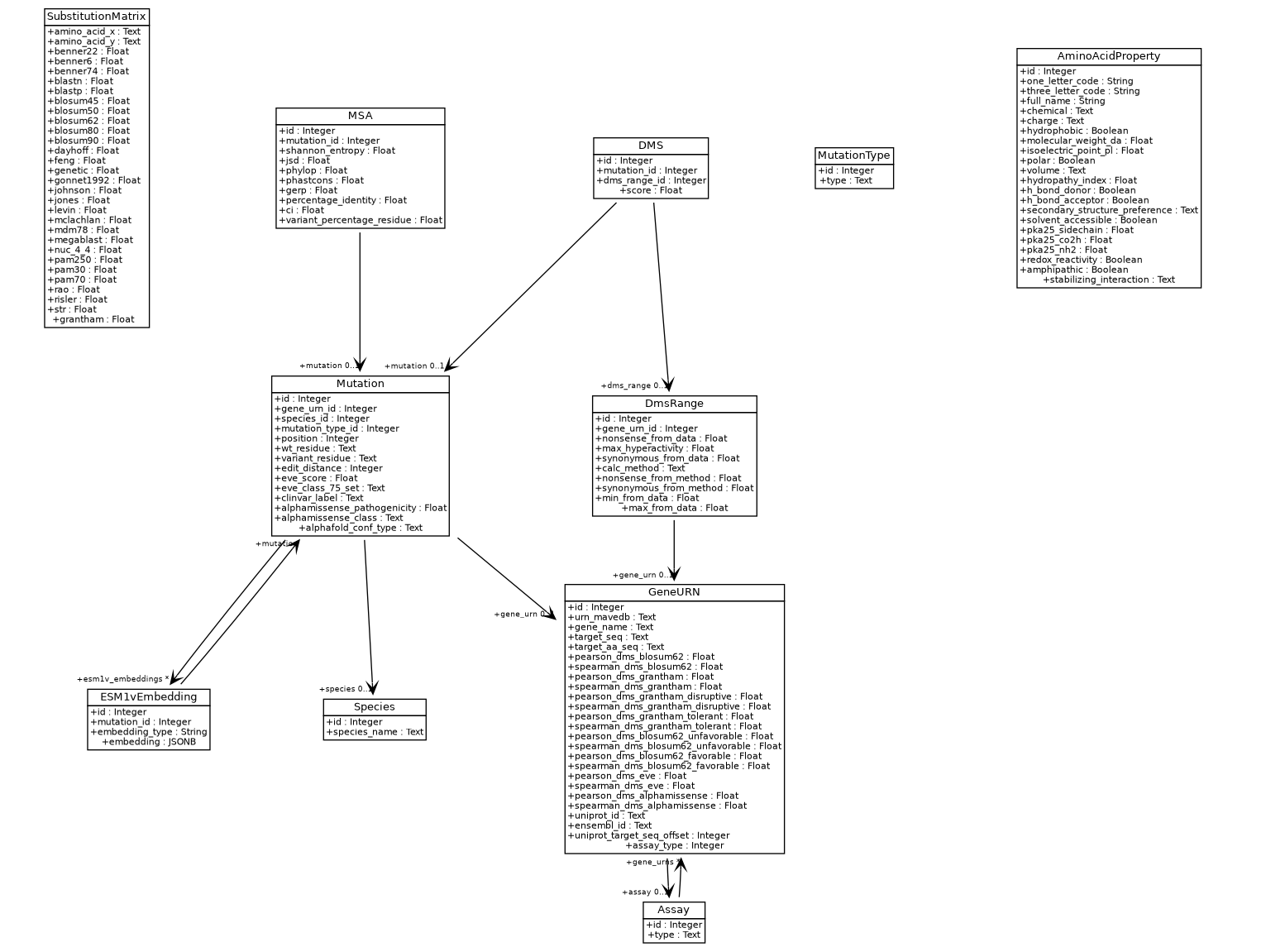

### Supplementary_Fig_S2.png

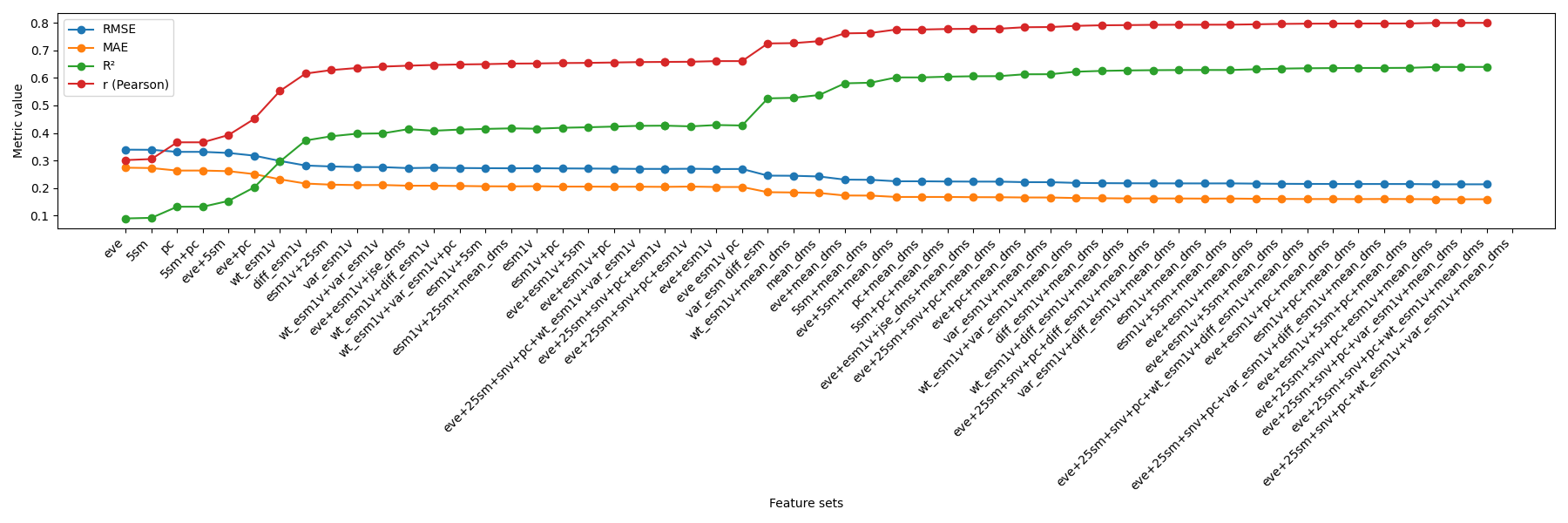

### Supplementary_Fig_S3.png

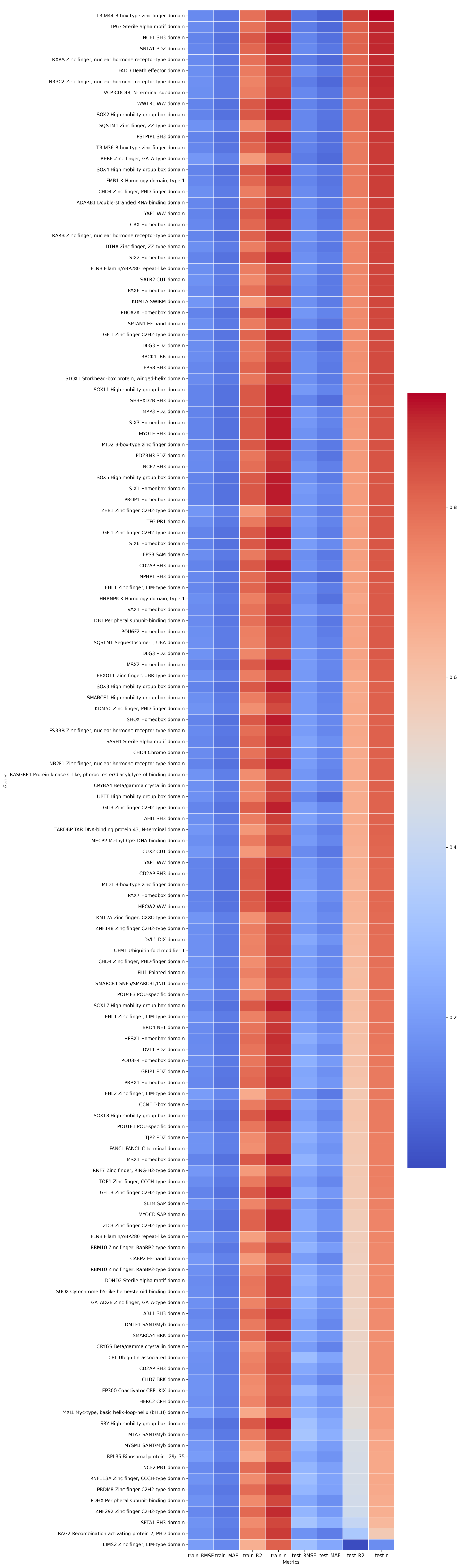

### Supplementary_Fig_S4.png

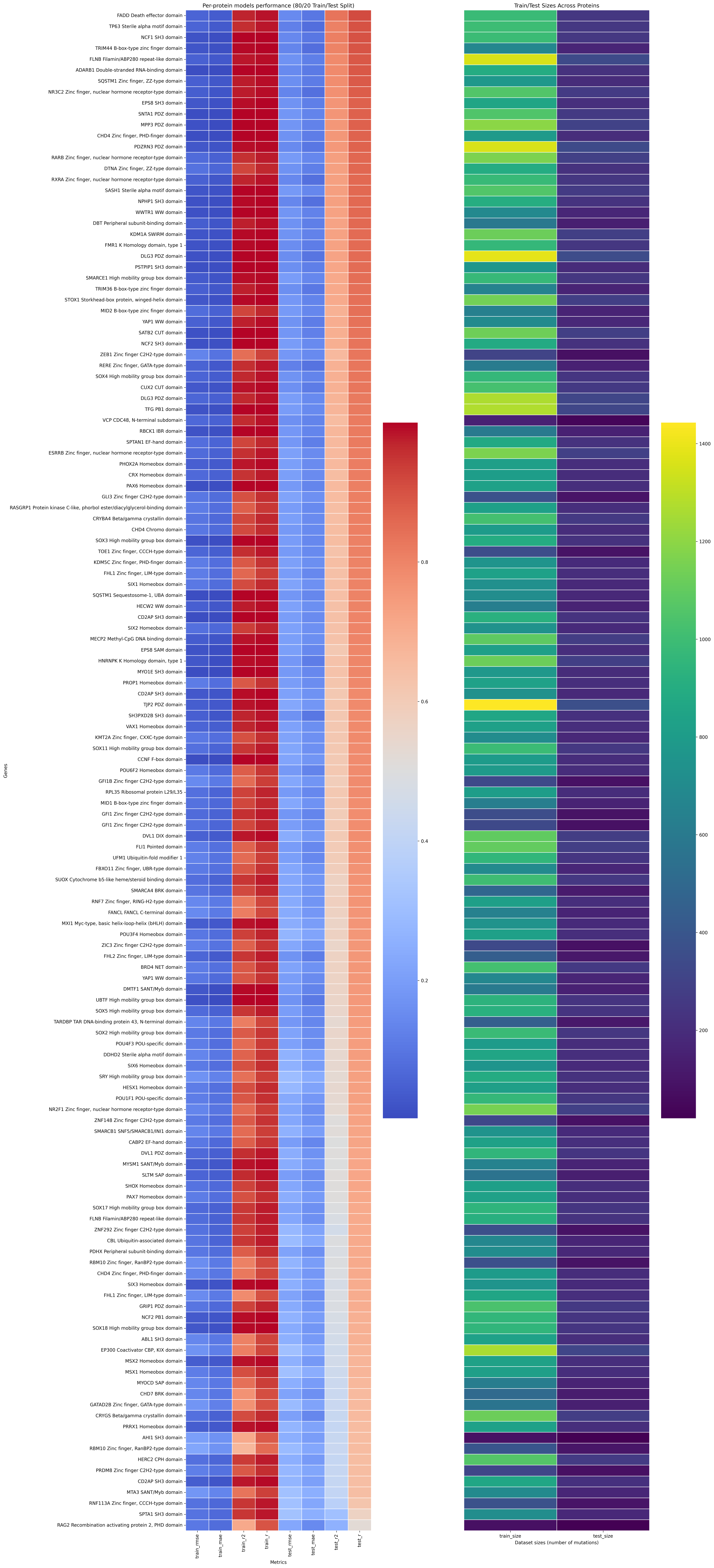

### Supplementary_Fig_S5.png

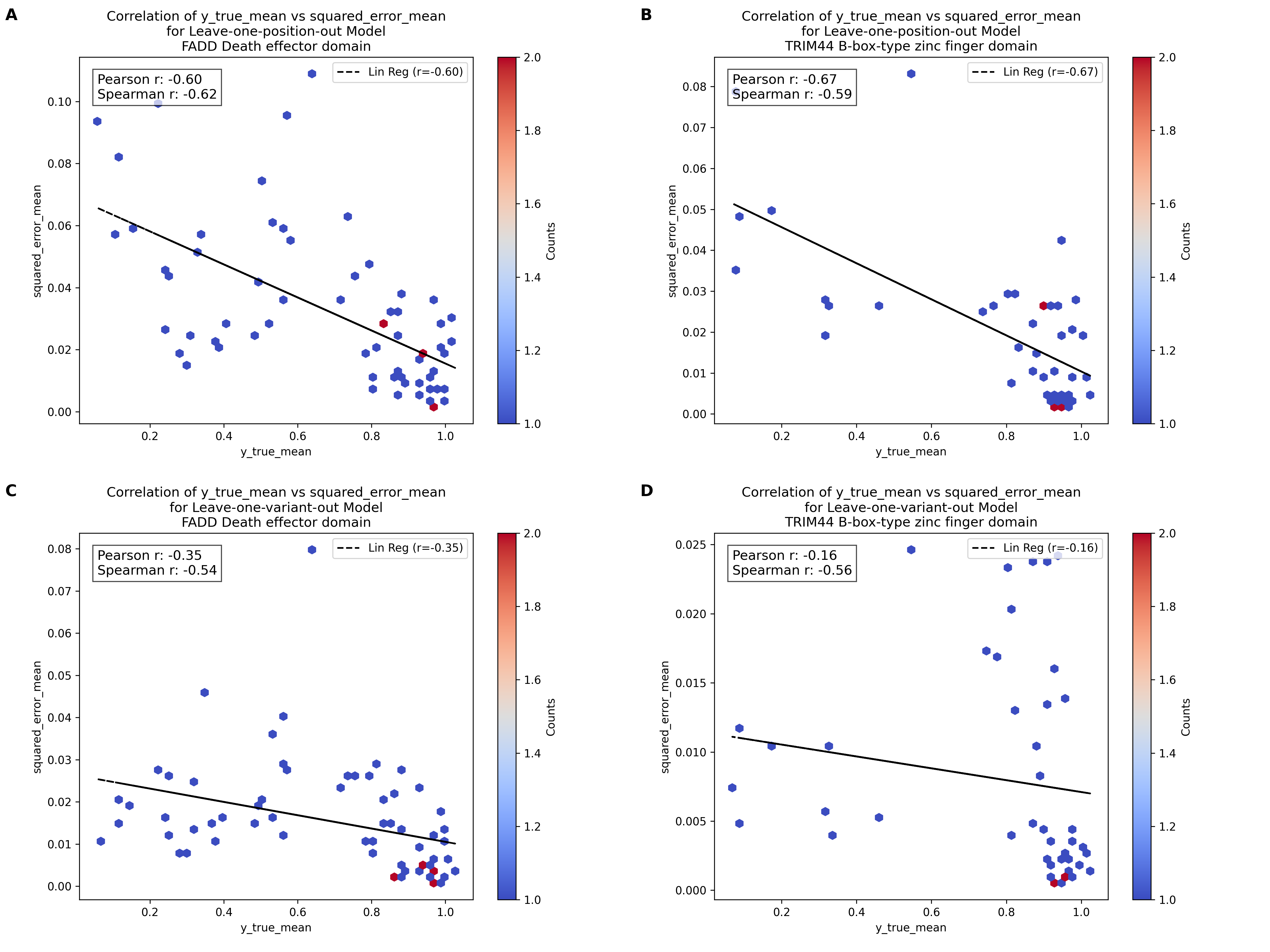

### Supplementary_Fig_S6.png

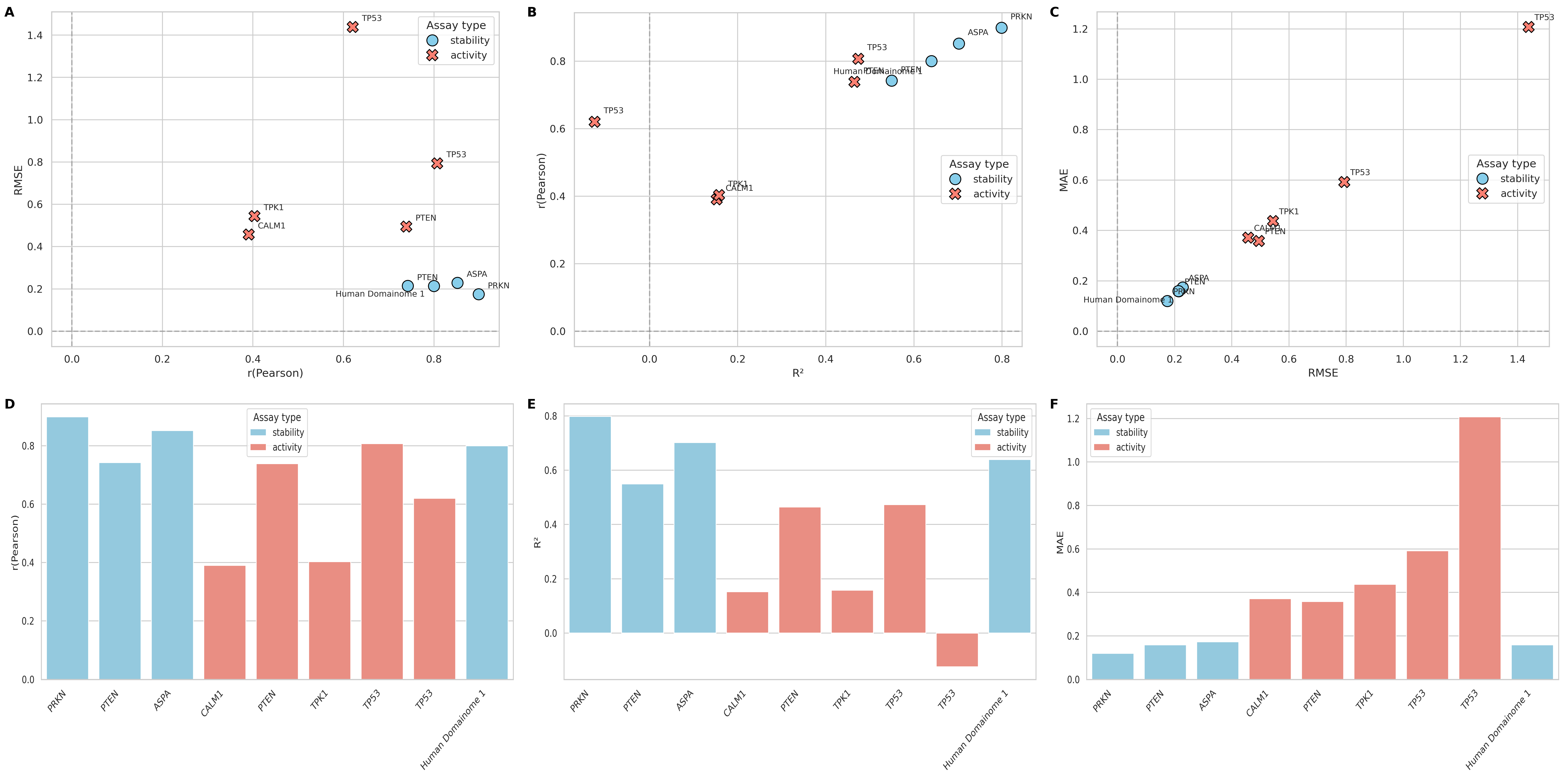

### Supplementary_Fig_S7.png

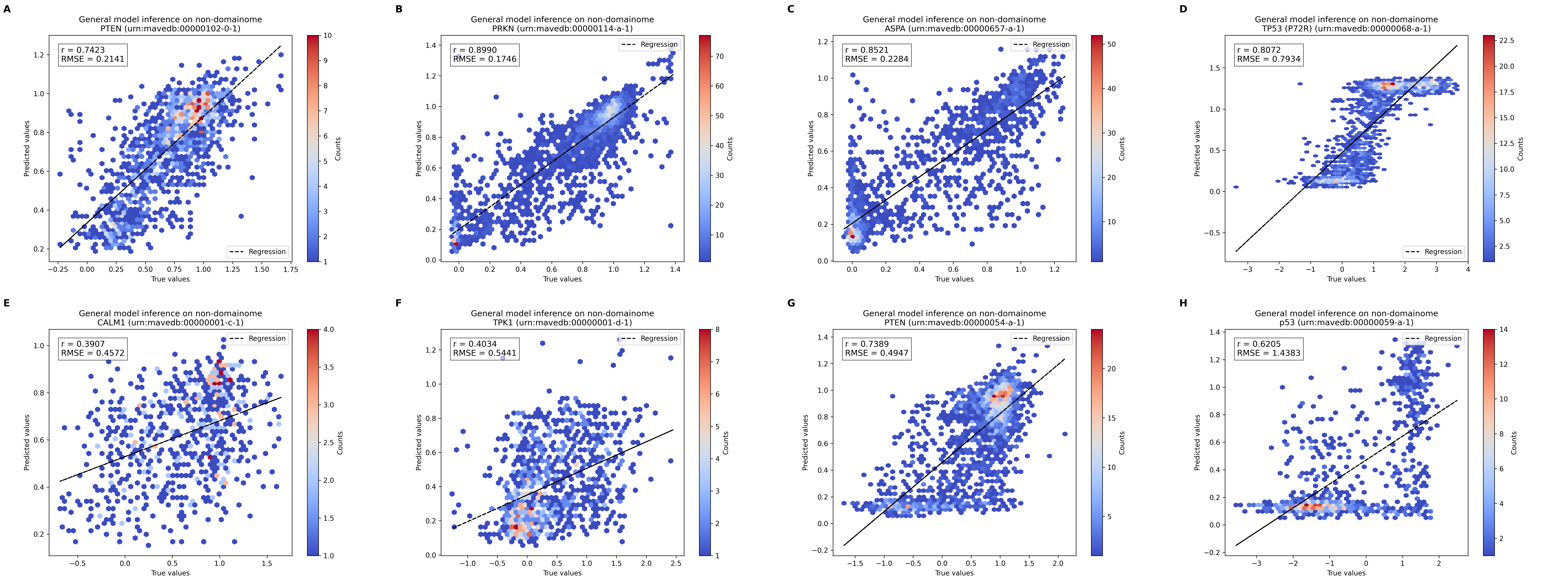

### Supplementary_Fig_S8.png

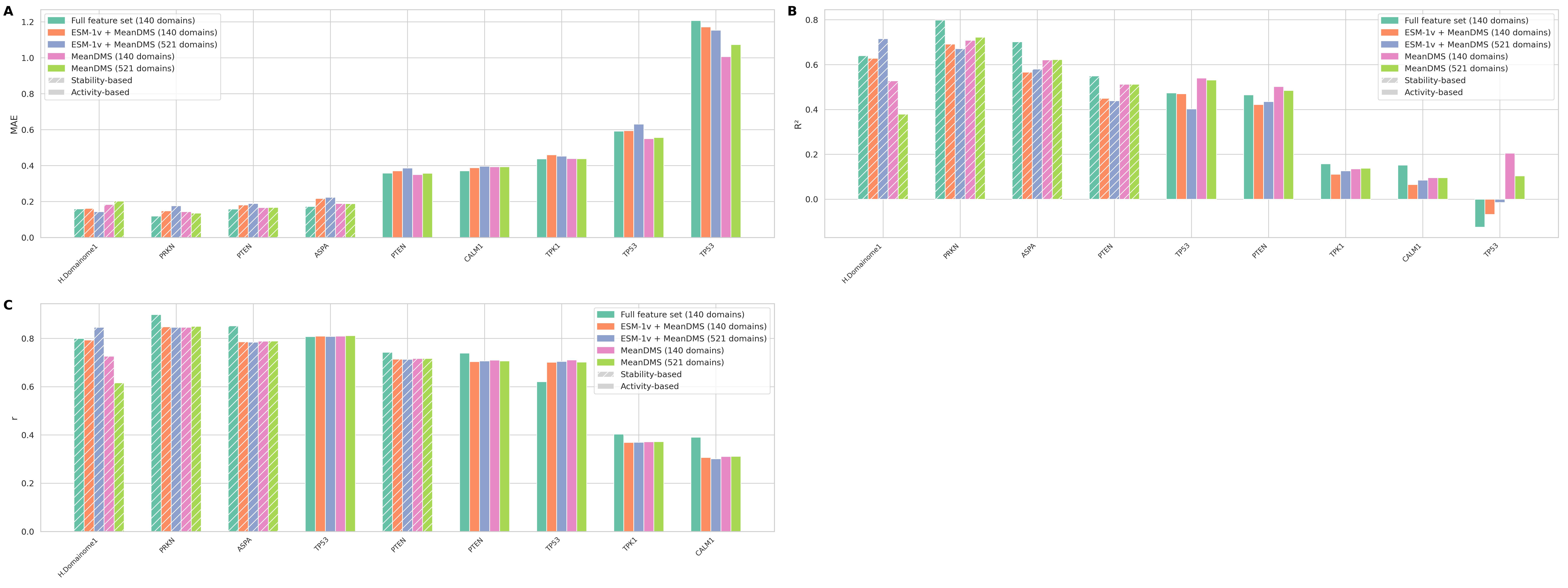

### Supplementary_Fig_S9.png

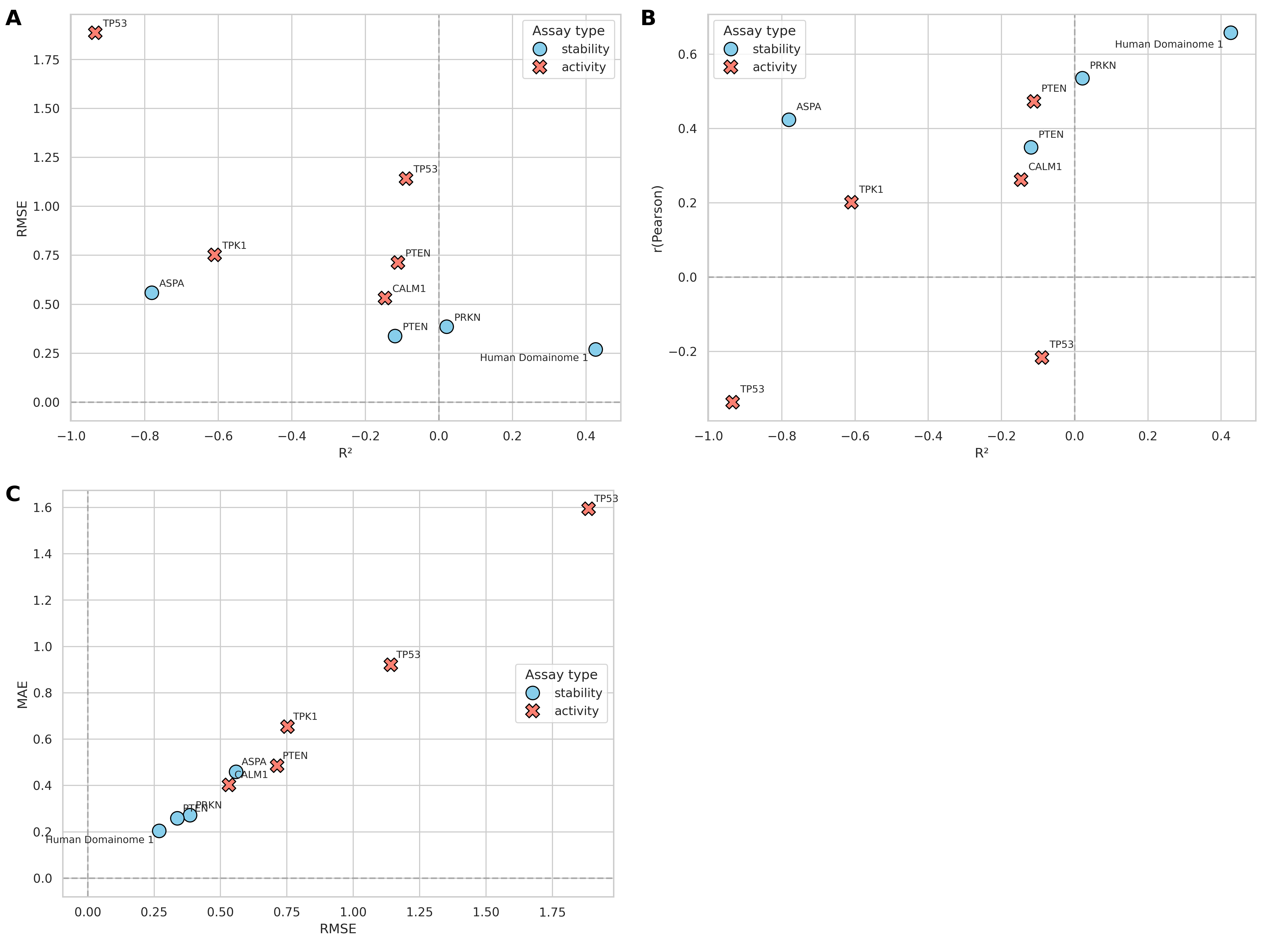

### Supplementary_Fig_S10.png

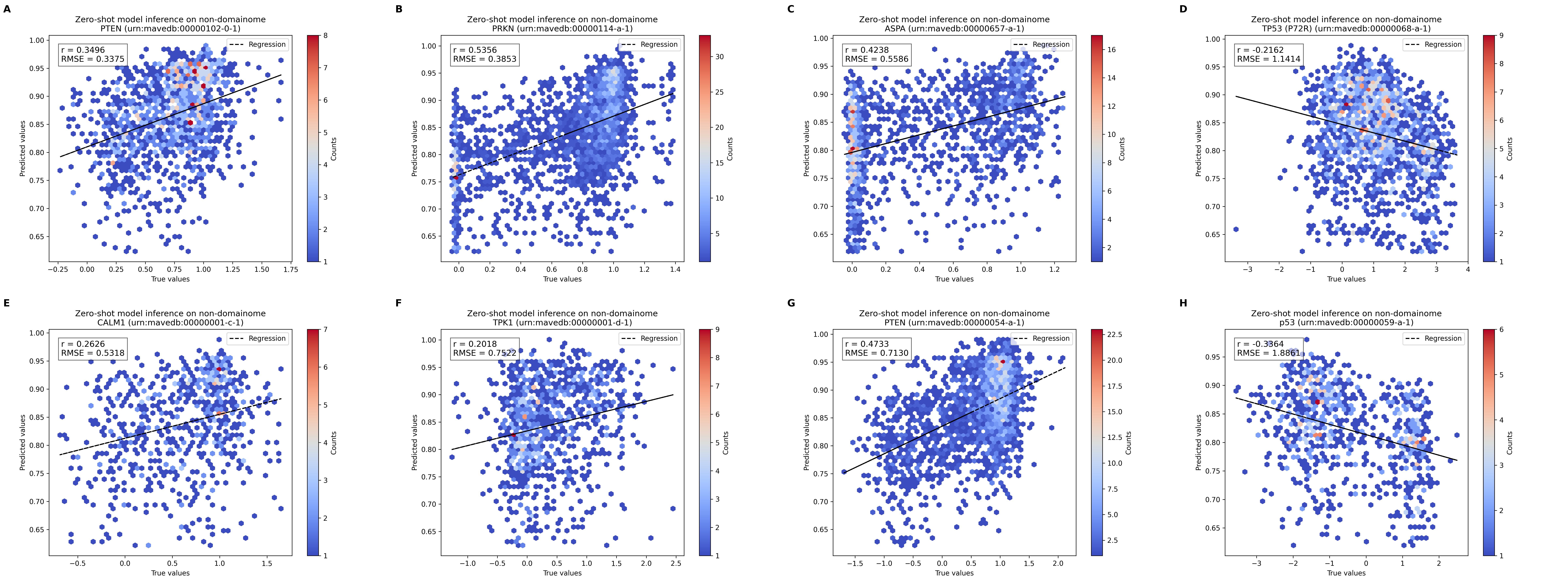
